# Distribution-free complex hypothesis testing for single-cell RNA-seq differential expression analysis

**DOI:** 10.1101/2021.05.21.445165

**Authors:** Marine Gauthier, Denis Agniel, Rodolphe Thiébaut, Boris P. Hejblum

## Abstract

State-of-the-art methods for single-cell RNA sequencing (scRNA-seq) Differential Expression Analysis (DEA) often rely on strong distributional assumptions that are difficult to verify in practice. Furthermore, while the increasing complexity of clinical and biological single-cell studies calls for greater tool versatility, the majority of existing methods only tackle the comparison between two conditions. We propose a novel, distribution-free, and flexible approach to DEA for single-cell RNA-seq data. This new method, called ccdf, tests the association of each gene expression with one or many variables of interest (that can be either continuous or discrete), while potentially adjusting for additional covariates. To test such complex hypotheses, `ccdf` uses a conditional independence test relying on the conditional cumulative distribution function, estimated through multiple regressions. We provide the asymptotic distribution of the `ccdf` test statistic as well as a permutation test (when the number of observed cells is not sufficiently large). `ccdf` substantially expands the possibilities for scRNA-seq DEA studies: it obtains good statistical performance in various simulation scenarios considering complex experimental designs (*i.e.* beyond the two condition comparison), while retaining competitive performance with state-of-the-art methods in a two-condition benchmark. We apply `ccdf` to a large publicly available scRNA-seq dataset of 84,140 SARS-CoV-2 reactive CD8+ T cells, in order to identify the diffentially expressed genes across 3 groups of COVID-19 severity (mild, hospitalized, and ICU) while accounting for seven different cellular subpopulations.

## 1. Introduction

Single-cell RNA sequencing (scRNA-seq) makes it possible to simultaneously measure gene expression levels at the resolution of single cells, allowing a refined definition of cell types and states across hundreds or even thousands of cells at once. Single-cell technology significantly improves on bulk RNA-sequencing, which measures the average expression of a set of cells, mixing the information in the composition of cell types with different expression profiles. New biological questions such as detection of different cell types or cellular response heterogeneity can be explored thanks to scRNA-seq, broadening our comprehension of the features of a cell within its microenvironment (Eberwine *and others*, 2014).

Several challenges arise from the sequencing of the genetic material of individual cells like in transcriptomics (see Lähnemann *and others* (2020) for a thorough and detailed review). Differential expression analysis (DEA) is a major field of exploration to better understand the mechanisms of action involved in cellular behavior. A gene is called differentially expressed (DE) if its expression is significantly associated with the variations of a factor of interest. Single-cell data have different features from bulk RNA-seq data that require special consideration for developing DEA tools. Indeed, scRNA-seq data display large proportions of observed zeros (i.e. “dropouts”), due either to biological processes or technical limitations (Lähnemann *and others*, 2020).

The large amount of single-cell measurements provides an opportunity to estimate and characterize the distribution of each gene expression and to compare it under different conditions in order to identify DE genes. In fact, scRNA-seq distributions usually show complex patterns. Therefore, the `scDD` method (Korthauer *and others*, 2016) condenses the difference in distribution between two conditions into four categories: the usual difference in mean, the difference in modality, the difference in proportions and the difference in both mean and modality. Since scRNA-seq data analysis lay unique challenges, new statistical methodologies are needed.

Several strategies making strong distributional assumptions on the data have been proposed to perform single-cell DEA. MAST (Finak *and others*, 2015) and SCDE (Kharchenko *and others*, 2014) are two well-know differential methods, the former using a two-part generalized linear model to take into account both the dropouts and the non-zero values by making a Gaussian assumption of each gene and the latter relying on a Bayesian framework combined with a mixture of Poisson and negative binomial distributions. `scDD` (Korthauer *and others*, 2016) makes use of a Bayesian modeling framework to detect differential distributions and then to classify the gene into four differential patterns using Gaussian mixtures. `DEsingle` (Miao *and others*, 2018) proposes a zero-inflated negative binomial (`ZINB`) regression followed by likelihood-ratio test to compare two samples. `D`^3^`E` (Delmans and Hemberg, 2016) also applies a likelihood-ratio test after fitting a Poisson-Beta distribution.

Yet, Risso *and others* (2018) have advanced that scRNA-seq data are zero-inflated and have proposed to use zero-inflated negative binomial models, while Svensson (2020) have argued that the number of zero values is consistent with usual count distributions. Then, Choi *and others* (2020) illustrated that scRNA-seq data are zero-inflated for some genes but “this does not necessarily imply the existence of an independent zero-generating process such as technical dropout”. In fact, biological information (e.g. cell type and sex) may explain it. The authors also discourage imputation as zeros can contain relevant information about the genes. In addition, Townes *and others* (2019) argue that single-cell zero inflation actually comes from normalization and log-transformation. The distribution and the sparsity of scRNA-seq data remains difficult to model, and – as there is no consensus on which model is the best one – it is of utmost importance to develop general and flexible methods for analyzing scRNA-seq data which do not require strong parametric assumptions.

Fewer distribution-free tools have been developed to model single-cell complex distributions without making any parametric assumption. EMDomics (Nabavi *and others*, 2016) and more recently SigEMD (Wang and Nabavi, 2018) are two non-parametric methods based on the Wassertein distance between two histograms, the latter including data imputation to handle the problem of the great number of zero counts. D^3^E offers in addition the possibility to perform either the Cramer-von Mises test or the Kolmogorov-Smirnov test to compare the expression values’ distributions of each gene, thus delivering a distribution-free option. In a comparative review, Wang *and others* (2019) illustrated that non-parametric methods, i.e. distribution-free, perform better in distinguishing the four differential distributions. Recently, Tiberi *and others* (2020) presented `distinct`, a hierarchical non-parametric permutation approach using empirical cumulative distribution functions comparisons. The method requires biological replicates (at least 2 samples per group) and allows adjustment for covariates but only tackles the comparison between two conditions to our knowledge.

However, the limitations of these state-of-the-art methods for scRNA-seq DEA are many. The approaches based on a distributional assumption face methodological issues. They rely on strong distributional assumptions that are difficult to test in practice. A deviation from the hypothesized distribution will translate into erroneous *p*-values and may lead to inaccurate results. While the increasing complexity of clinical and biological studies calls for greater tools versatility, the majority of existing methods, whether parametric or non-parametric, cannot handle data sets with a complex design, making them very restrictive. In fact, the most commonly used methods remain in the traditional framework of DEA restricted to the comparison between two conditions only. One might be interested in the genes differentially expressed across several conditions (e.g. more than two cell groups, multi-arm...) or in testing the genes differentially expressed according to a continuous variable (e.g. cell surface markers measured by flow cytometry...). In particular, the identification of surrogate biomarkers from transcriptomic measurements is becoming an emerging field of interest, especially in cancer therapy (Wang *and others*, 2007) or in new immunotherapeutic vaccines (Cliff *and others*, 2004). Gene expression could be used to compare treatments in observational settings. Yet in such cases, adjusting for some technical covariates or some confounding factors is paramount to ensure the validity of an analysis, as this external influence can impact the outcome as well as the dependent variables and thus generates spurious results by suggesting a non-existent link between variables.

Overall, the need of testing the association between gene expression and the variables of interest, potentially adjusted for covariates, makes an additional motivation for developing suitable tools. The complex hypothesis we aim to test consists in performing a conditional independence test (CIT). A CIT broadens the classical independence test by testing for independence between two variables given a third one, or a set of additional variables (see Figure 8). Two random variables *X* and *Y* are conditionally independent given a third variable *Z* if, and only if, *P* (*X, Y* | *Z*) = *P* (*X* | *Z*)*P* (*Y* | *Z*). As described in Li and Fan (2020), many CIT have been developed previously and are readily available such as discretization-based tests (Huang *and others*, 2010), metric-based tests (Runge, 2018; Su and White, 2007; Huang *and others*, 2016), permutation-based two-sample tests (Doran *and others*, 2014; Sen *and others*, 2017), kernel-based tests (Muandet *and others*, 2017; Li *and others*, 2009) and regression-based tests (see Li and Fan (2020) for a short review). Yet, these CIT either suffer from the curse of dimensionality or are hardly applicable to a large number of observations (Muandet *and others*, 2017). Zhou *and others* (2020) have converted the conditional independence test into an unconditional independence test and then used the Blum–Kiefer–Rosenblatt correlation (Blum *and others*, 1961) to develop an asymptotic test. Yet, the latter cannot be applied when *X* is discrete. Those limitations make these tests impractical in our context of scRNA-seq DEA and thus require adaptation.

Performing DEA necessarily involves performing as many independent tests as there are genes. The variables of interest may be either discrete or continuous, while the number of covariates to condition upon may also increase. Consequently there is an urgent need for a CIT that is both flexible and fast. Here, we propose a novel, distribution-free, and flexible approach, called `ccdf`, to test the association of gene expression to one or several variables of interest (continuous or discrete) potentially adjusted for additional covariates. Because of the current limitations of existing CIT and the growing interest in testing differences in distribution, we make use of a CIT based on the conditional cumulative distribution function (CCDF), estimated by a regression technique. We derive the asymptotic distribution of the `ccdf` test statistic, which does not rely on the underlying distribution of the data, as well as a permutation test to ensure a good control of type I error and FDR, even with a limited sample size.

Section 2 describes our proposed method with both asymptotic and permutation tests. Section 3 presents several simulation scenarios to illustrate the good performances of `ccdf` when we consider complex designs (*i.e.* beyond the two condition comparison), while a benchmark in the two condition case shows our method retains similar performance in terms of statistical power compared to competitive state-of-the-art methods. Section 4 compares the performances of our method with several methods on a “positive data set” that included differentially expressed genes as well as a “negative data set” without any differentially expressed gene. Section 5 illustrates the application of `ccdf` to a scRNA-seq study in COVID-19 patients. Section 6 discusses the strengths and limitations of our proposed approach.

## 2. Method

In this section, we propose a new, easy-to-use, fast, and flexible test for scRNA-seq DEA. We give both its asymptotic distribution as well as a permutation approach to obtain valid *p*-values in small samples.

### 2.1 Conditional independence test

#### 2.1.1 Null hypothesis

Testing the association of *Y*, namely the expression of a gene, with a factor or a group of factors of interest *X* either discrete (multiple comparisons) or continuous given covariates *Z* is equivalent to test conditional independence between *Y* and *X* knowing *Z*: “ *H*_0_ : *Y* ⊥ *X* | *Z* “. Several statistical tools can be used to characterize the probability law of a random variable, such as the characteristic function, the probability density distribution or the cumulative distribution function. The characteristic function is not often used in practice, due to its relative analytical complexity. The probability density distribution, while more popular, remains difficult to estimate in practice when the number of variables increases due to the increasing number of bandwidths to be optimized. This curse of dimensionality quickly leads to classical computational problems because of complexity growth. As for the cumulative distribution function, its estimation does not require any parameter akin to these bandwidths, making it an efficient tool in highdimensional non-parametric statistics. From this point, we built a general DEA method including an estimation of the CCDF based on regression technique.

The conditional independence test we propose is based on CCDFs. Indeed, if a group of factors is associated with the expression of a gene, the immediate consequence is that the CCDF of the gene expression would be significantly different from the marginal cumulative distribution, which overlooks this conditioning. Thus, the null hypothesis can be written as: “ *H*_0_ : *F_Y_* _|*X,Z*_ (*y, x, z*) = *F_Y_* _|*Z*_ (*y, z*) “, where the CCDF of *Y* given *X* and *Z* is defined as *F_Y_* _|*X,Z*_ (*y, x, z*) = ℙ(*Y* ≤ *y* | *X* = *x, Z* = *z*). If there are no covariates, the conditional independence test turns into a traditional independence test as we test the null hypothesis *Y* ⊥ *X* which is equivalent to test *F_Y_* _|*X*_ (*y, x*) = *F_Y_* (*y*).

#### 2.1.2 Test statistic

In this section, we propose a general test statistic for testing the null hypothesis of conditional independence that is easy to compute. We denote 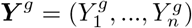 an outcome vector (i.e. normalized read counts for gene *g* in *n* cells) and 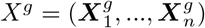 a *s* × *n* matrix encoding the condition(s) to be tested that can be either continuous or discrete. One may want to add exogenous variables, which are not to be tested but upon which it is necessary to adjust the model. Let 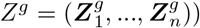 be a *r* × *n* matrix for continuous or discrete covariates to take into account. For the sake of simplicity, we drop the notation *g* in the remainder as we refer to gene-wise DEA.

We have ***Y*** ∈ [*ζ*_min_*, ζ*_max_] for some known constants *ζ_min_, ζ_max_*. Let *ζ*_min_ ≤ *ω*_1_ *< ω*_2_ < … < *ω_p_ < ζ*_max_ be a sequence of *p* ordered and regular thresholds. For each *ω_j_* with *j* = 1, …*, p*, the CCDF *F_Y_* _|_ *X,Z* (*ω*_*j* | *x, z*_) may be written as a conditional expectation 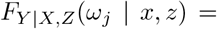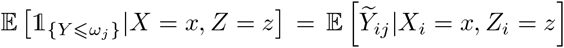 where 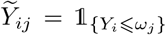 is a binary random variable that is 1 if *Y_i_* ≤ *ω_j_* and 0 otherwise. We propose to estimate these conditional expectations through a sequence of *p* working models:

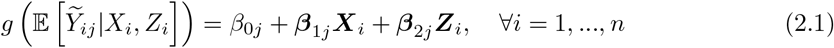

where 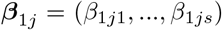 is the vector of size *s* referring to the regression of 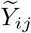 onto ***X**_i_* and 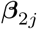 is the vector of size *r* referring to the regression of 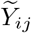 onto ***Z**_i_*, for the fixed thresholds *ω*_1_*, ω*_2_, …, *ω_p_*. If *X* has no link with *Y* given *Z*, then we expect that 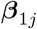 will be null. So, we aim to test:

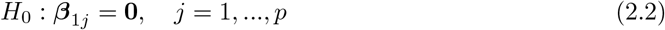

Although, there are many different test statistics associated with this null hypothesis, we propose to use the following test statistic that can be written as 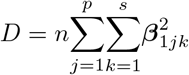.

#### 2.1.3 Estimation and asymptotic distribution

In this section, we describe how to estimate 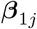, which allows the computation of the test statistic *D*.

While in principle any link function *g*(·) could be selected for the models (2.1), we select the identity link *g*(*y*) = *y* for its computational simplicity, and we compute coefficient estimates using ordinary least squares (OLS). Because our approach requires *p* models for each of possibly thousands of genes, speed is of utmost importance, so we use OLS. Other selections of *g*(·) are of course possible and could be explored at the cost of additional computation time.

We show in the Supplementary Material that using OLS to estimate (2.1), 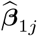 can be expressed by 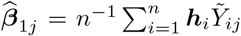, where ***h***_*i*_ is a function of the design matrix *W* with *i*th row ***W**_i_* = (1*, **X**_i_, **Z**_i_*). These estimates may be plugged in to obtain the estimated test statistic 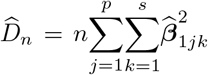. We furthermore show in the Supplementary Material that the asymptotic distribution of the test statistic may be approximated by a mixture of 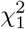 random variables:

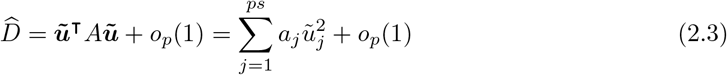

where 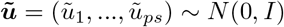 are standard multivariate normal random variables and *a_j_* are the eigenvalues of 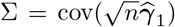 where 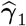 is a vectorized version of 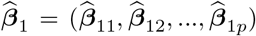 (the *s* × *p* matrix) concatenating the *s* rows of 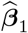 one after another.

We may then compute *p*-values by comparing the observed test statistic 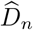 to the distribution of 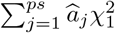, where 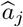 is an estimate of *a_j_* based on a consistent estimator for Σ (see Supplementary Material for details). Therefore, this allows us to derive a *p*-value for the significance of a gene with regards to the variable(s) to be tested. In practice, we make use of saddlepoint approximation for distributions of quadratic forms (Kuonen, 1999; Chen and Lumley, 2019) to compute *p*-values for the mixture of *χ*^2^s, implemented in the `survey` R package (Lumley, 2004). Note that we obtain a simple limiting distribution without relying on any distributional assumptions on the gene expression. In fact, based on the results in Li and Duan (1989), our test will be asymptotically valid as long as there exist any *g*(·) and any 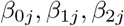 such that (2.1) holds. Lastly, the great number of tests requires the Benjamini–Hochberg (Benjamini and Hochberg, 1995) correction afterwards, which is automatically applied to the raw *p*-values in the R package of `ccdf`.

#### 2.1.4 Permutation test

Permutation tests are recommended when the number of observations is too small, so that the asymptotic distribution can not be assumed to hold. When the sample size *n* is low, we propose to perform permutations to estimate the empirical distribution of 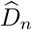 under the null hypothesis. The test is described in the Supplementary Material, along with practical considerations for computational speed up.

## 3. Simulation study

### 3.1 Comparisons with state-of-the-art methods in the two conditions case

We compared the performance of our method `ccdf` with three state-of-the-art methods, `MAST`, `scDD` and `SigEMD`, to find differentially distributed (DD) genes. We have selected methods implemented in R that have shown good results in Wang *and others* (2019) benchmarks and that have been especially designed for single-cell data (excluding methods for bulk RNA-seq like `edgeR` and `DESeq2`). Also, we considered methods that use normalized (and therefore continuous) counts as input, so that the results of our simulations are comparable. We generated simulated count data from negative binomial distributions and mixtures of negative binomial distributions. Since `MAST`, `scDD` and `SigEMD` require continuous input, we transformed the counts into continuous values while preserving as much as possible the count nature of the data (*i.e.* negative binomial assumption). To do so, we added a small Gaussian noise with mean equal to 0 and variance equal to 0.01 to the simulated counts. 500 simulated datasets were generated including 10,000 genes each, of which 1,000 are differentially expressed under a two conditions setting for 7 different sample sizes *n* (20, 40, 60, 80, 100, 160, 200). The observations were equally divided into the two groups. Korthauer *and others* (2016) classified four different patterns of unimodal or multi-modal distributions: differential expression in mean (DE), differential proportion (DP), differential modality (DM) and both differential modality and different component means within each condition (DB). We used these four differential distributions to create our own simulations. Specifically, we simulated 250 DD genes, 250 DM genes, 250 DP genes and 250 DB genes. Plus, 9,000 non-differentially expressed genes are simulated according to two non-differential scenarios (see Supplementary Materials for the simulation settings).

Figure 2 shows the Monte-Carlo estimation over the 500 simulations of the type-I error and the statistical power as well as the false discovery rate and true discovery rate, according to increasing samples sizes. The type-I error is computed as the proportion of significant genes among the true negative and the power as the proportion of significant genes among the true positive. After Benjamini-Hochberg correction for multiple testing, the FDR is computed as the proportion of false positives among the genes declared DD and the TDR as the proportion of true positives among the genes declared DD. The *p*-value nominal threshold is fixed to 5%. The three state-of-the-art methods as well as `ccdf` exhibit good control of type-I error and no inflation of FDR. The four methods present a high overall True Discovery Rate. However, `MAST` shows a lack of power of about 23% for a sample size equal to 200. The True Positive rate (after Benjamini-Hochberg correction) for each scenario for all the methods is shown Figure 3. The three leading methods perform well in finding the traditional difference in mean (DE) as soon as a number of 60 observations is reached. `ccdf` is less powerful for a sample size of 20 and 40 cells but shows the same performances with a larger sample size. The difference in modality (DM) is well detected by all the methods with a slight advantage for `SigEMD` in low sample sizes (from 20 to 60 cells). The difference in proportion (DP) is not favorable for the asymptotic test of `ccdf` until 160 observations but the permutation test exhibits higher power. The asymptotic test requiring a sufficiently large number of observations to converge, the permutation test is more efficient for a lower number of cells. `SigEMD` and `scDD` are the most effective in detecting DP genes. Even though `MAST` shows good power for DE, DM and DP genes, it fails to detect DB genes. In fact, `MAST` is designed to detect difference in the overall mean (traditional differential expression), which is absent in DB scenario. The difference in modality and in different component means is then overlooked as expected with its parametric model. `SigEMD`, `scDD` and both of `ccdf`’s tests present competitive performances. Generally speaking, `ccdf` retains competitive performance with the state-of-the-art in this two condition benchmark, which makes it a method particularly adapted to traditional DEA. Even though `ccdf` is a non-parametric method, the asymptotic test is relatively reasonable in terms of computation times, especially compared to `SigEMD` (see Table 1). If the computation times seem too large, we recommend to use the adaptive thresholds strategy explained in the Supplementary Materials and in particular for the permutation test, we recommend to switch to the adaptive permutations also described in the Supplementary Materials.

**Fig. 1.**
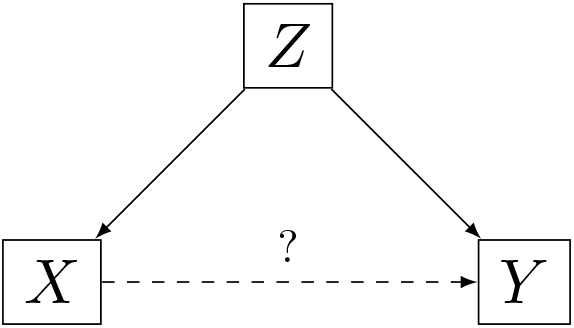
Conditional dependence graph (Li and Fan, 2020)

**Fig. 2.**
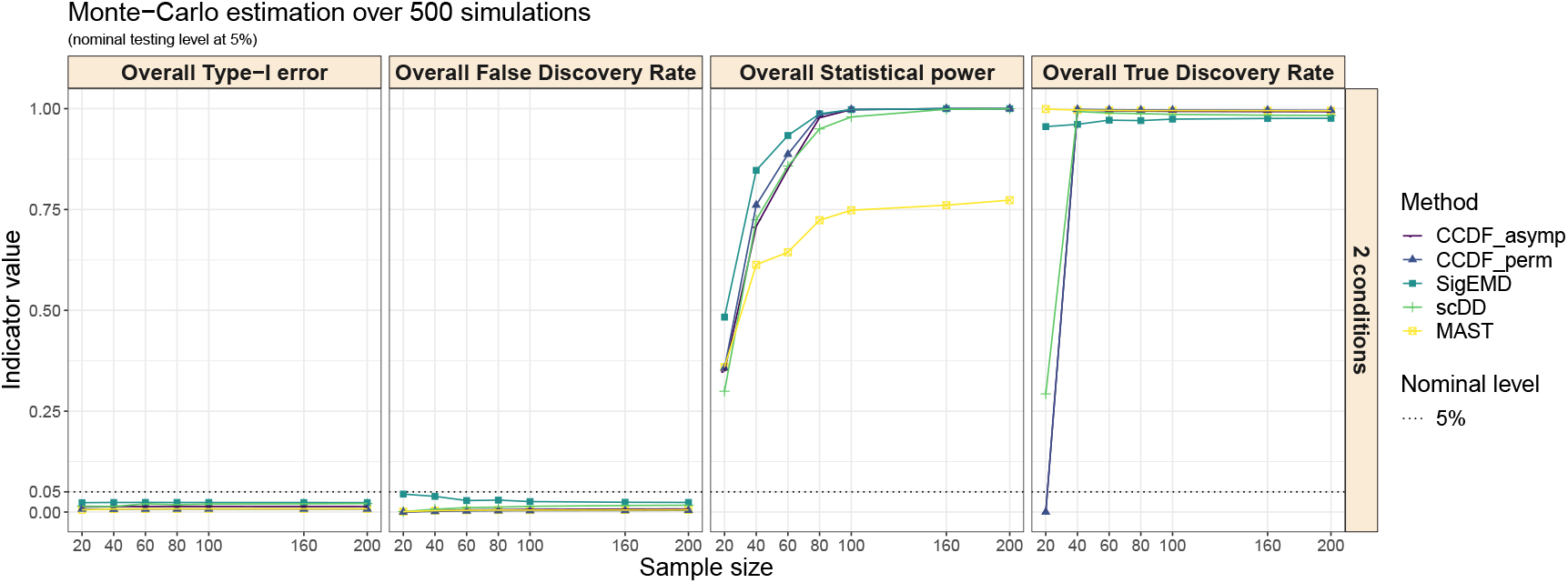
Overall Type-I error, Power, FDR and TDR under the 2 conditions case with increasing sample size. For `ccdf`, we perform the asymptotic test and the permutation test.

**Fig. 3.**
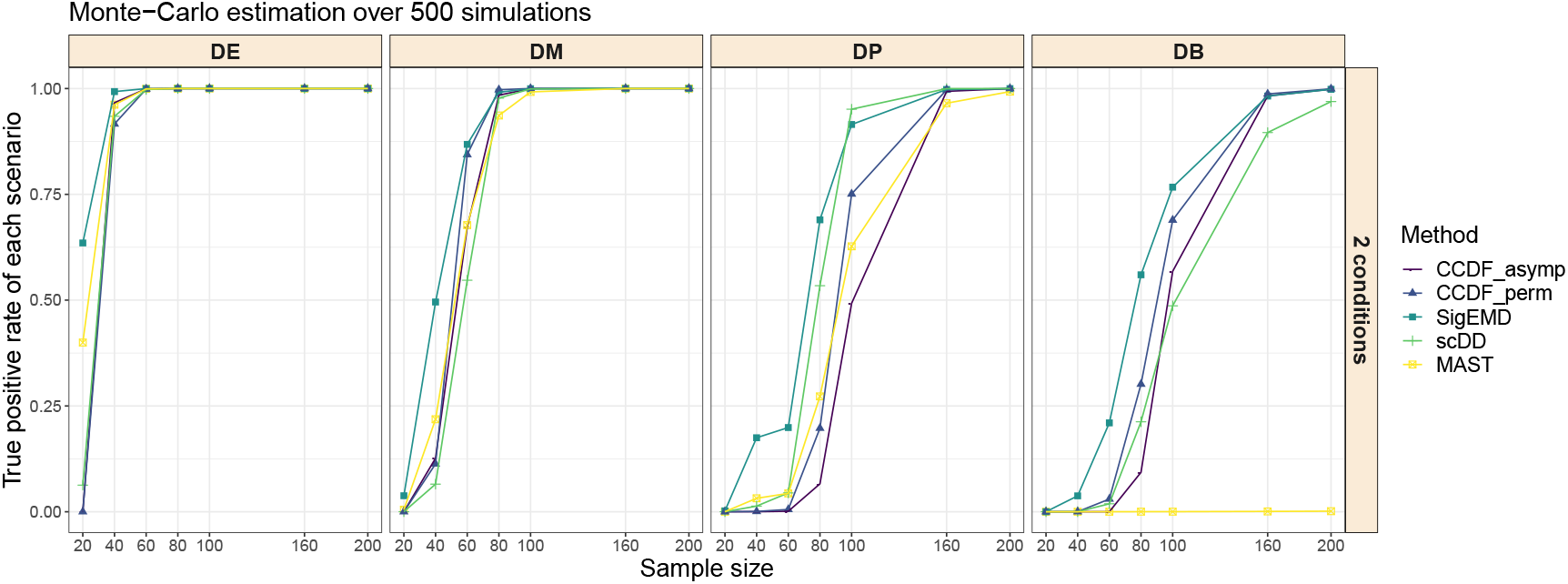
True positive rate under the 2 conditions case for the four DD scenarios with increasing sample size. DE: difference expression in mean. DM: difference in modality. DP: difference in proportion. DB: both differential modality and different component means within each condition. For `ccdf`, we perform the asymptotic test and the permutation test.

**Table 1.** Computation times for the state-of-the-art methods and `ccdf` in the two condition case for one run and *n* = 100, using a MacBook Pro 2019 (2,3 GHz Intel Core i9 8 cores 16 threads)

| Method | Computation times in minutes |
| --- | --- |
| ccdf asymptotic test | 3 |
| ccdf permutation test | 1385 |
| ccdf permutation test with adaptive permutations | 750 |
| SigEMD | 1680 |
| MAST | 1.4 |
| scDD | 0.05 |

### 3.2 Multiple comparisons

This second scenario deals with a multiple comparisons design where cell observations were split into 4 different groups. Count data were generated from negative binomial distributions and mixtures of negative binomial distributions and transformed into continuous values as in section 3.1. 500 simulated datasets were generated including 10,000 genes each, of which 1,000 are differentially expressed under a four conditions setting for 7 different sample sizes *n* (40, 80, 120, 160, 200, 320, 400). The observations were then equally divided into four groups. Instead of generating two distributions for each gene as in section 3.1, we created four distributions and therefore new DD scenarios for this specific DEA simulation: multiple DE, multiple DP, multiple DM and multiple DB (more details in the Supplementary Materials). The non-differentially genes are also simulated in a specific fashion described in the Supplementary Material. The idea of differential distribution patterns was converted into a multiple differential distribution setting. For example, the multiple DD scenario consists in four distributions with four different means. Since the other approaches can not handle this type of design, only `ccdf` with the asymptotic test and the permutation test is run. Figure 5 shows that both versions of `ccdf` have great power to identify DE and DM genes as the sample size increases. DP and DB genes require larger number of cells to achieve sufficient power (from *n* > 200).

**Fig. 4.**
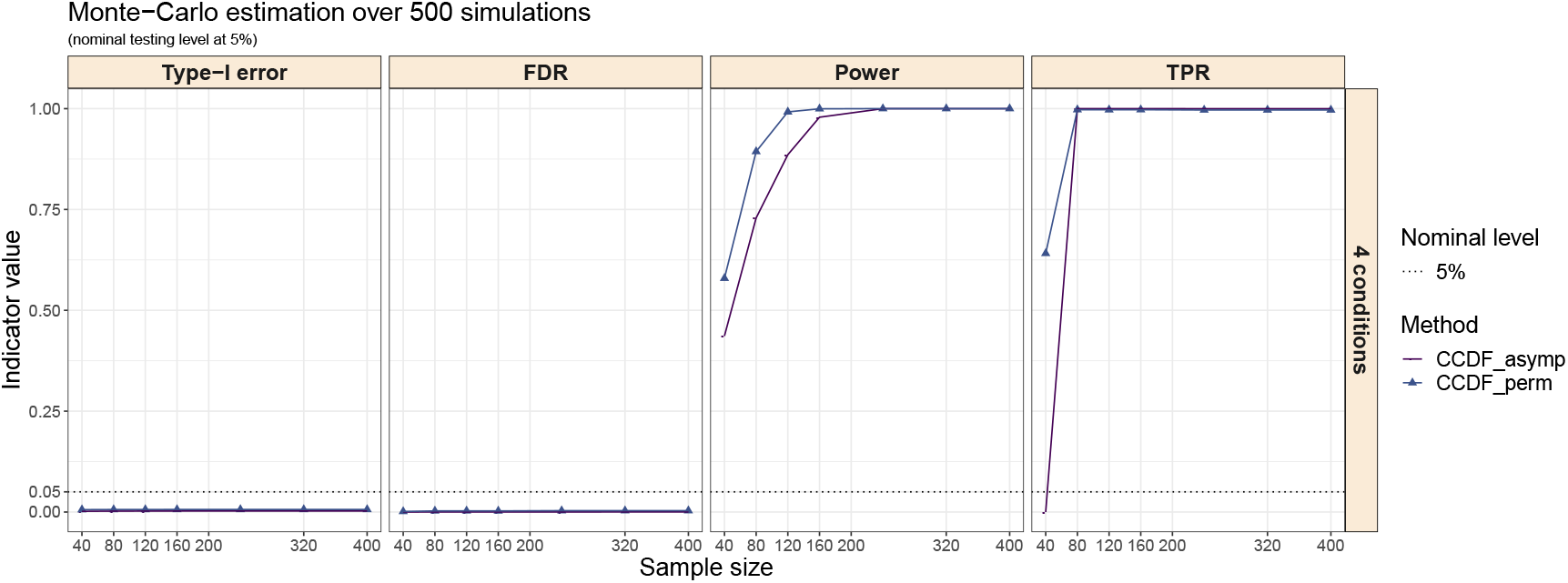
Overall Type-I error, Power, FDR and TDR for ccdf under the 4 conditions case with increasing sample size. `ccdf` is the only method capable of dealing with more than 2 conditions. We perform the asymptotic test and the permutation test.

**Fig. 5.**
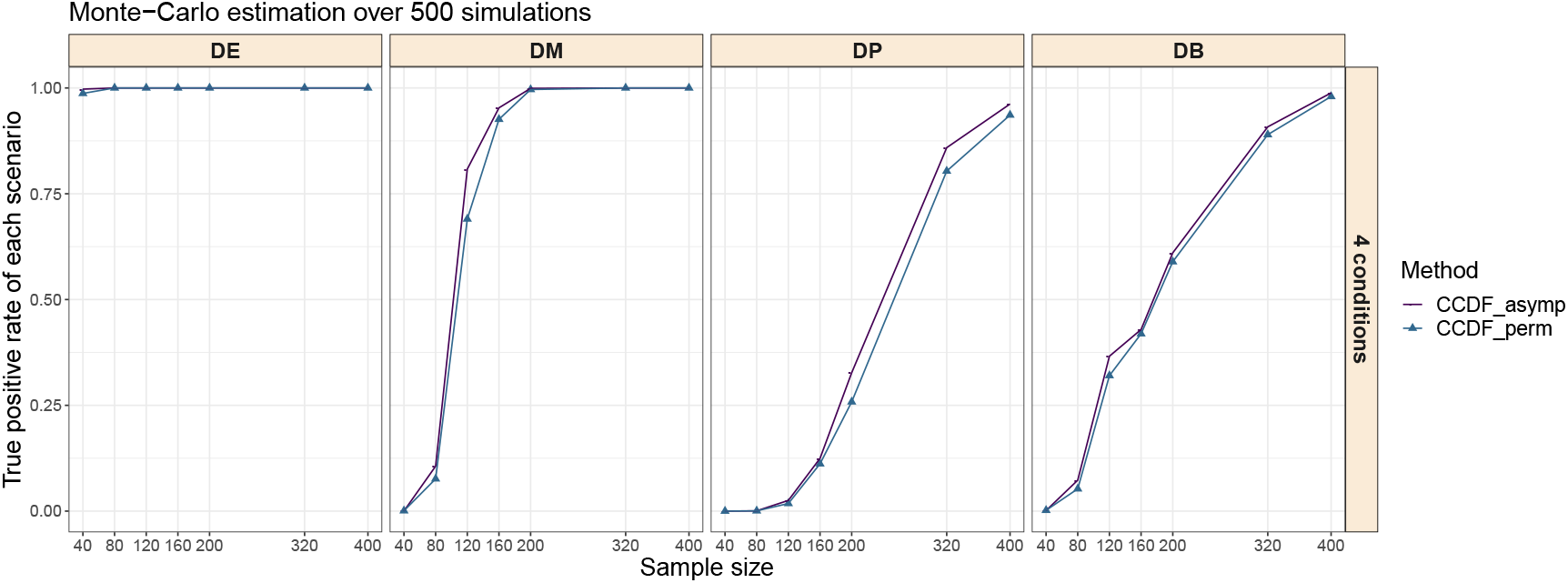
True positive rate under the 4 conditions case for the four DD scenarios with increasing sample size. DE: difference expression in mean. DM: difference in modality. DP: difference in proportion. DB: both differential modality and different component means within each condition.`ccdf` is the only method capable of dealing with more than 2 conditions. We perform the asymptotic test and the permutation test.

### 3.3 Two conditions comparison given a covariate Z

As represented in Figure 8, a potential confounding covariate *Z* can interfere the test of the link between the outcome *Y* and the variable *X*. Then, it is needed to adjust for the confounding variable by carrying out a CIT. We aim to emphasize the importance of taking into account *Z*, thanks to our approach, by showing the erroneous results obtained by not doing so. For this purpose, we simulated a confounding variable *Z* from a Normal distribution. The values of the variable to be tested *X* depends on the quartile of *Z*, creating a strong link between *X* and *Z*. *Y* is constructed from *X* for DE genes and from *Z* for non-DE genes, the last case is particularly misleading if one does not take into account *Z* (see simulation settings in Supplementary Material).

We simulated 10,000 genes of which 1,000 are differentially expressed for several sample sizes n (20, 40, 60, 80, 100, 160, 200). `ccdf` permutation test was excluded in this simulation scheme because of the large amount of time to compute. In fact, when we have to adjust for a covariate, the permutation test is not suited to such large sample sizes and number of genes because of the underlying permutation strategy.

`MAST` is able to adjust for covariates like `ccdf` so we expect good performances from both of them. Conversely, `scDD` and `SigEMD` can not control for confounding variables that might impact the detected DE genes. The results of the benchmark between `MAST`, `scDD`, `SigEMD` and `ccdf` are depicted in Figure 6. Under the alternative hypothesis, we created a large difference in the mean of the normal distributions between the two conditions in order to make DE genes easier to identify. We see therefore that `scDD`, `SigEMD` and `ccdf` exhibit high power at all sample sizes whereas `MAST` tends to be less powerful for a given size. As expected, the link between *Y* and *Z* is very confusing for `scDD`, `SigEMD` which interpret it as a link between *Y* and *X*, since *X* is constructed from *Z*. Consequently, we observe a consistent rise of Type-I error. `ccdf` and `MAST` perfectly control the Type-I error. In addition, `scDD` and `SigEMD` suffer greatly in terms of TDR as the number of cells is increasing. In fact, fewer real discoveries are found meaning that more genes are identified as significant while being actually false positive, which is in line with the drastically inflated FDR. Dealing with this complex design, `ccdf` outperforms the leading methods by controlling the Type-I error as well as the FDR and by providing a powerful test. This simulation study highlights the importance of taking into account the confounding variables that may exist. Otherwise, DEA may lead to inaccurate results and a potential huge amount of false positives. It is worth mentioning that `ccdf` can adjust for more than one covariate using the asymptotic test while preserving relatively fast calculation times. The permutation test is for now limited to one adjustment variable and the computation times are obviously increased due to the permutation strategy.

**Fig. 6.**
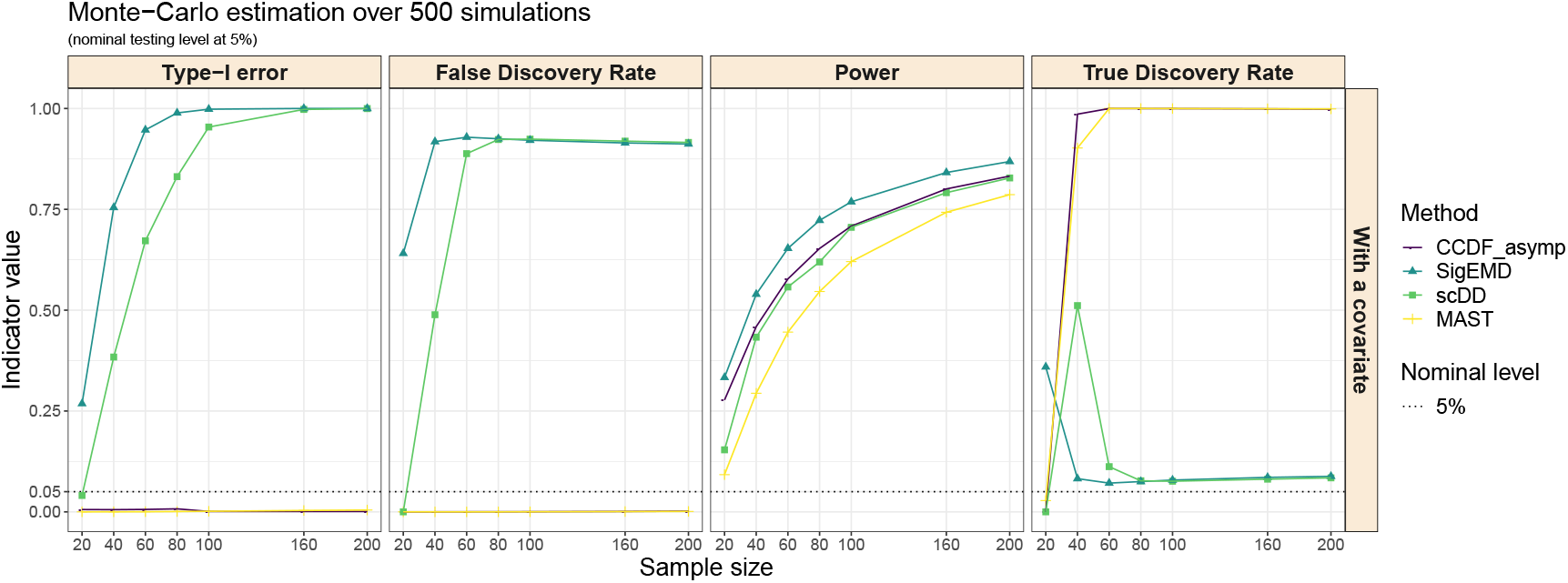
Overall Type-I error, Power, FDR and TDR under the 2 conditions comparison given a confounding variable with increasing sample size. For `ccdf`, we perform the asymptotic test and the permutation test.

## 4. Comparisons using real data benchmarks

To be in a more realistic context with a greater number of zeros, a positive control data set and a negative control data set described in Wang *and others* (2019) were used in order to compare the performances of several methods. The genes with a variance equal to zero were removed from the datasets and counts were converted into log-transformed count per millions (CPM) values.

### 4.1 Positive control dataset

The dataset from Islam *and others* (2011) includes 22,928 genes measured across 48 mouse embryonic stem cells and 44 mouse embryonic fibroblasts. We considered 1,000 genes validated through qRT-PCR experiments as a gold standard gene set in the same fashion as Wang *and others* (2019) in order to compute true positive rate. A gene is defined as a true positive if it is found as DE by the method and belongs to the gold standard set (Moliner *and others*, 2008). For `ccdf`, `SigEMD`, `scDD` and `MAST`, the number of DE genes for an adjusted *p*-value of 0.05 and the proportion of true positives among those identified as DE are given in Table 2. `ccdf` leads to the highest true positive rate (0.734) and enables to identify 24.6% more genes in common with the top 1,000 genes compared to `SigEMD` (0.488). `scDD` shows a true positive rate of 0.351 and `MAST` exhibits the lowest rate (0.198) of all the tools.

**Table 2.** Number of detected DE genes, and sensitivities of the state-of-the-art methods and `ccdf` tools using positive control real data for an adjusted *p*-value of 0.05

| Method | Number of DE genes | (TP/1000 gold standard) |
| --- | --- | --- |
| *ccdf* | 9,353 | 0.734 |
| SigEMD<sup>†</sup> | 3,702 | 0.488 |
| scDD<sup>†</sup> | 2,638 | 0.351 |
| MAST<sup>†</sup> | 734 | 0.198 |

<sup>†</sup> results from Wang *and others* (2019).

### 4.2 Negative control dataset

To get false positive rate, we used the dataset from Grün *and others* (2014). We selected 80 samples under the same condition. To create 10 datasets, we randomly divided these 80 cells into 2 groups of 40 cells. As there is no difference between the two groups, not a single DE gene is to be found in each dataset. Performance evaluations are compared across methods in Table 3. `ccdf` and `MAST` do not detect any genes which was expected. Although all the cells are under the same condition, `scDD` and `SigEMD` identified respectively 5 and 50 DE genes.

**Table 3.** Number of the detected DE genes and false positive rates of the state-of-the-art methods and `ccdf` using negative control real data for an adjusted *p*-value of 0.05

| Method | Number of detected DE genes | False positive rate |
| --- | --- | --- |
| `ccdf` | 0 | 0 |
| `scDD`<sup>†</sup> | 5 | 0.0007 |
| `MAST`<sup>†</sup> | 0 | 0 |
| `SigEMD`<sup>†</sup> | 50 | 0.007 |

<sup>†</sup> results from Wang *and others* (2019).

**Table 4.** Study design

|  | Healthy subjects | COVID-19 patients |  |  |
| --- | --- | --- | --- | --- |
|  |  | Not admitted | non-ICU (hospitalized) | ICU (hospitalized) |
| **Sex** |  |  |  |  |
| Male | 7 | 7 | 10 | 7 |
| Female | 1 | 9 | 3 | 2 |
| **Total** | 8 | 16 | 13 | 9 |

**Table 5.** `Seurat` clustering

|  | Cluster |  |  |  |  |  |  |  |
| --- | --- | --- | --- | --- | --- | --- | --- | --- |
|  | 0 | 1 | 2 | 3 | 4 | 5 | 6 | 7 |
| **Number of cells** | 27086 | 17915 | 12270 | 9783 | 5688 | 5170 | 829 | 156 |

## 5. Application to a scRNA-seq study in COVID-19 patients

Kusnadi *and others* (2021) publicly released a large scRNA-seq data set on COVID-19, which is available from GEO (GSE153931). It includes UMI counts for 13,816 genes measured across 84,140 virus-reactive CD8+ T cells. Cellular gene expression was measured in 38 COVID-19 patients and 8 healthy controls. Among the COVID-19 patients, 9 were treated in Intensive Care Unit (ICU), 13 were treated in standard hospital wards (non-ICU) and 16 were not hospitalized at all. This data set features a particularly complex design for DEA (see table 8). Kusnadi *and others* (2021) regrouped the 22 hospitalized patients as having a severe COVID-19 while the 16 non-hospitalized were considered as mild, and they compared the gene expression across CD8+ T cells between them. Instead, we went further by analyzing the differences between the three COVID-19 categories, namely ICU, non-ICU hospitalized and not hospitalized to provide better insights between the different severity of COVID-19.

As is often done in scRNA-seq analyses, Kusnadi *and others* (2021) clustered the 84,140 cells, resulting in 8 different clusters. Table 8 presents the numbers of cells assigned to each cluster (Kusnadi *and others* (2021) excluded cluster 7 due to its small size). These clusters represent various CD8+ T cell sub-populations with different states of cell differentiation, with cluster 1 being composed of exhausted (dying) cells for instance. Some biological pathways, activated across some clusters, could also be associated to the disease severity (such as pro-survival features as reported in the original paper (Kusnadi *and others*, 2021)). Hence, it is not clear whether the association of the expression of some genes with the severity of the disease is either reflecting a difference in abundance of some clusters of CD8+ T cell populations, or due to the activation of some biological pathways independently of the clusters. We propose to compare the gene expression according to the severity of the disease while adjusting or not for the clusters, which might be acting as a confounding factor. Both the outcome and the potential confounding factor are categorical variables with more than 2 levels, making our analysis design relatively complex.

With our notations, *Y* is the gene expression, *X* the COVID-19 severity status and *Z* the cell cluster. We apply the asymptotic `ccdf` test for both analyses (with and without adjusting on *Z*) while testing each gene for differential expression across COVID-19 severity status. The number of thresholds to compute the marginal and conditional CDFs is set to 10. Following Kusnadi *and others* (2021), only transcripts expressed in at least 0.1% of the cells were included in the differential analyses, yielding 10,525 genes in total to be tested and UMI data are then converted to *log*_2_(CPM+1). 6,290 genes are found DE when adjusting for the clusters, while 8,181 genes are DE in unadjusted analyses (see Figure 7).

**Fig. 7.**
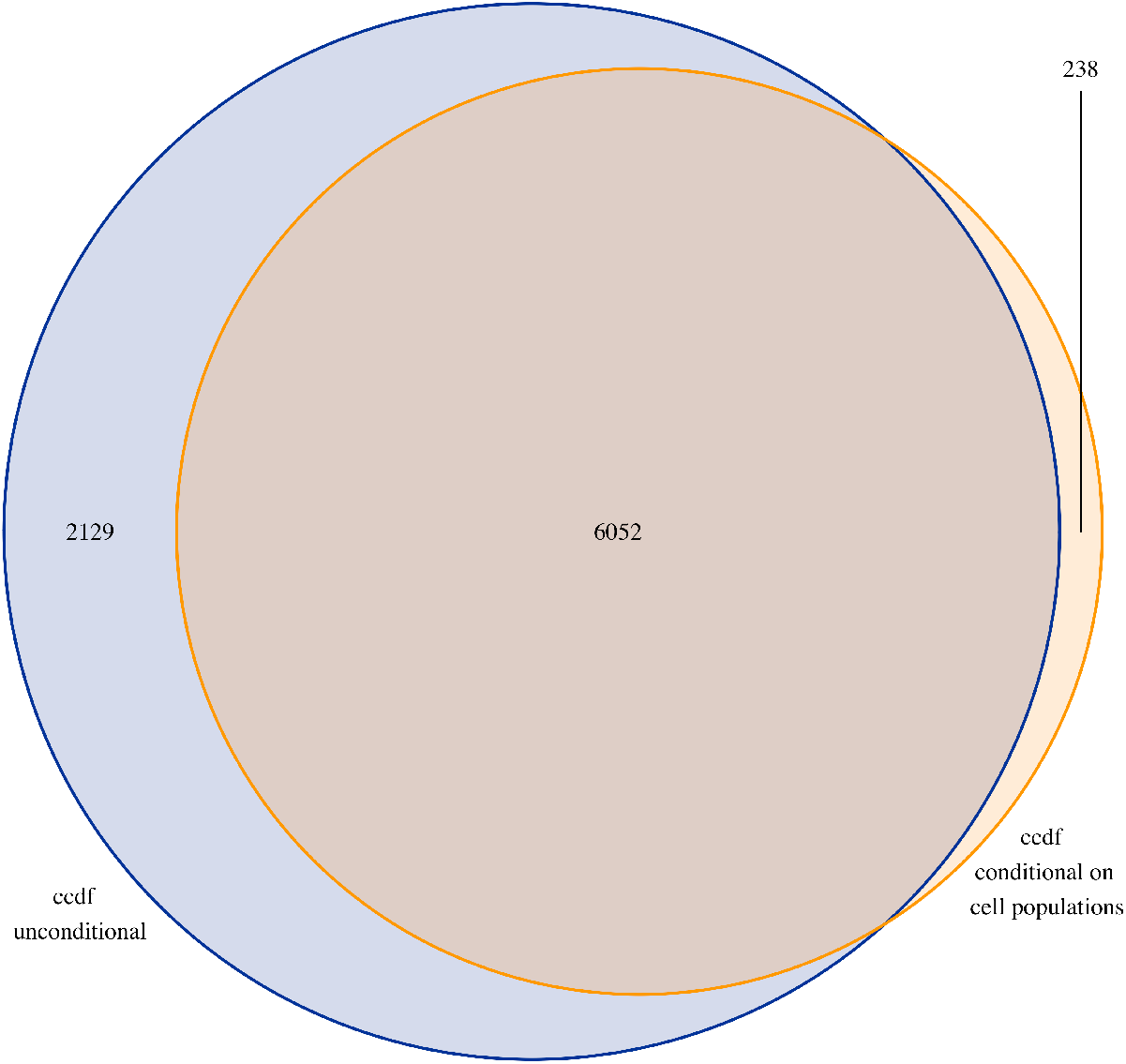
Venn diagram of the gene signature found by `ccdf` conditional on 7 cell populations and the gene signature found by `ccdf` without conditioning on the cell populations (`ccdf` unconditional)

**Fig. 8.**
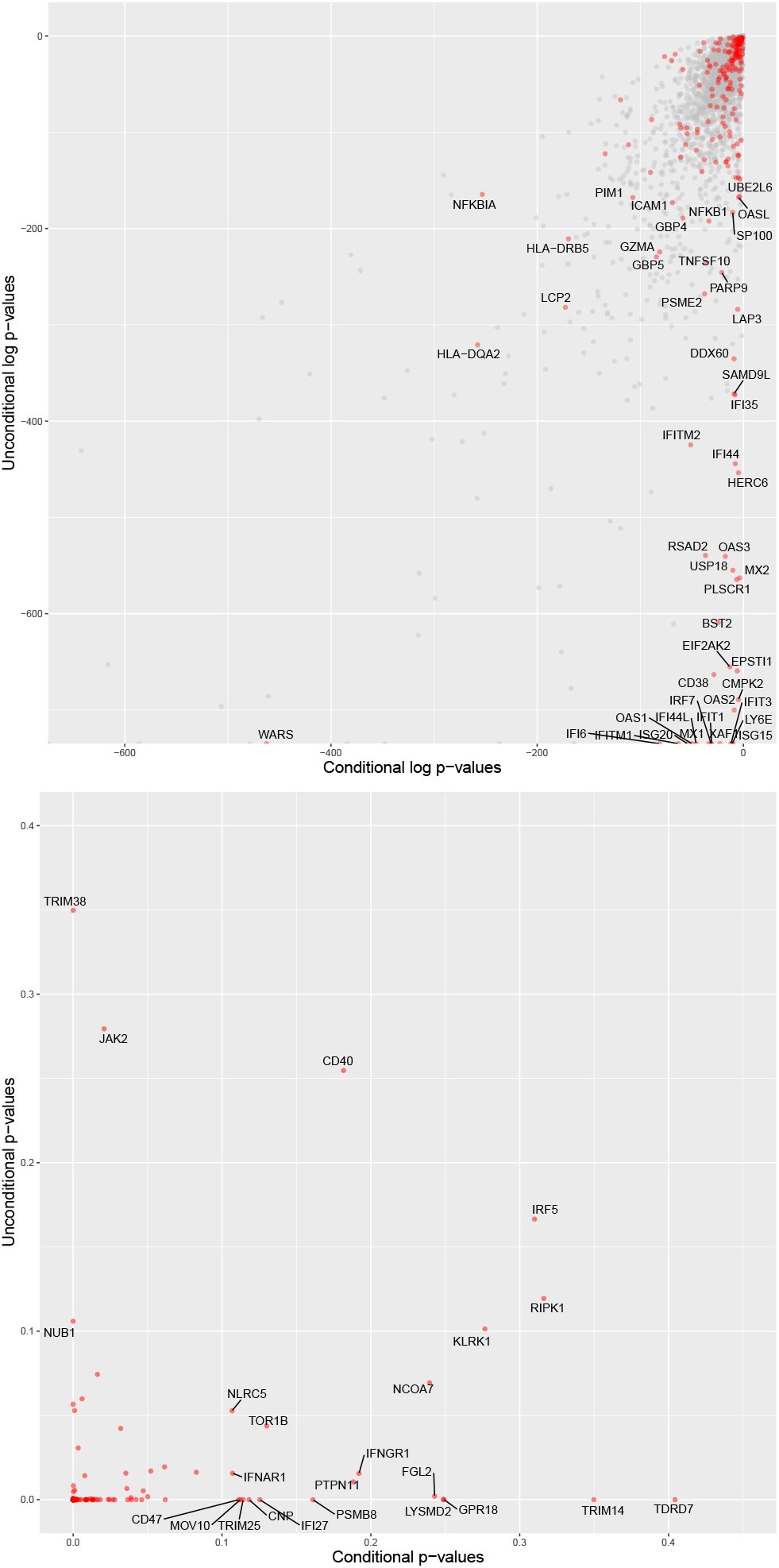
Top figure: scatter plot of the genes *p*-values in log-scale. The x-axis represents the log *p*-values obtained with `ccdf` conditional on cell populations. The y-axis represents the log *p*-values obtained with `ccdf` without conditioning on the cell populations. The red points correspond to the genes of the IFN list. The grey points correspond to the genes that are not part of the IFN list. Bottom figure: scatter plot of the IFN genes *p*-values between 0 and 0.05. The x-axis represents the *p*-values obtained with `ccdf` conditional on cell populations. The y-axis represents the *p*-values obtained with `ccdf` without conditioning on the cell populations.

To understand where the difference of 2,129 DE genes comes from, we focused specifically on the 221 genes included in the IFN gene set, one of the 7 pathways highlighted by Kusnadi *and others* (2021) (see their supplementary material Table S5 for their definition). Among these genes, 188 were found significant regardless of the adjustment on the clusters while 7 become significant and 20 become not significant when adjusting for clusters. Figure 8 shows that some genes with low *p*-values in unadjusted DEA have much higher *p*-values after adjustment for clusters (e.g. IFIT3, IFI6, TDRD7, IFI44L, IFI44, IFITM2, OAS2, HERC6 and CD38) whereas others such as LCP2 do not exhibit a strong variation of the *p*-value. This could be expected when looking at the abundance of the genes according to the clusters (Table S4 in Kusnadi *and others* (2021)) as for instance IFIT3, IFI6, IFI44 were much more abundant in cluster 1 than in other clusters, whereas LCP2 abundance was more balanced across clusters. As a visualization example, Figure 9 displays the various conditional CDFs estimated through both `ccdf` analyses for TDRD7 gene. It illustrates the reduction of the difference between the conditional CDF on *X* and its marginal counterpart (when not conditioning on *X*) when additionally conditioning on *Z* the clusters. This is reflected by the p-value which increase from 1.796244e-06 to 0.4043629 with the conditioning on *Z*: while TDRD7 remains DE in both analyses, its significance is decreased when taking into account the different cell clusters. While the majority of IFN genes remain significant at a threshold of 5% in both DEA, there is a general increasing trend of the *p*-values in adjusted analyses. This supports some confounding effect of the clusters on the significance of some genes within the Interferon pathway. Thus, taking into account the clusters does not fundamentally change the biological interpretation that Interferon pathway plays an important role in the severity status. But a part of the difference in abundance of some genes is due to the difference in abundance of the identified clusters. See supplementary figures for additional visualizations of `ccdf` results.

**Fig. 9.**
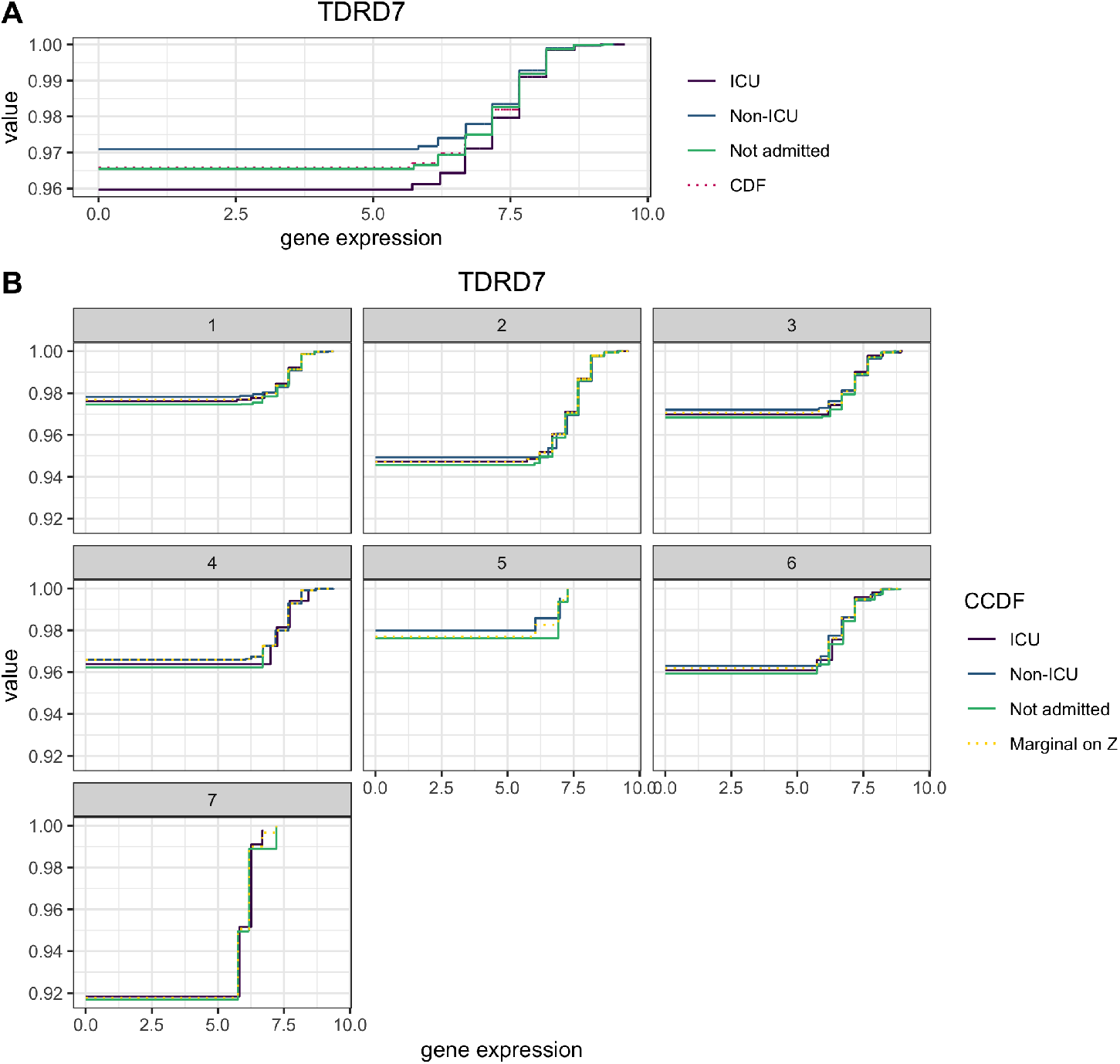
A: The solid lines represent the conditional CDF of TDRD7 gene on ICU status (ICU, Non-ICU and Not admitted) and the dark pink dotted line represents the marginal CDF of TDRD7 gene, *i.e.* without conditioning on ICU status. The underlying test performed by `ccdf` in the first DEA consists in comparing the marginal CDF with the conditional CDF. B: The solid lines represent the conditional CDF of TDRD7 gene on both *Z*, the 7 clusters, and *X*, the severity status (ICU, Non-ICU and Not Admitted), while the dotted yellow line represents the marginal CDF of TDRD7 expression without conditioning on *X* (but only conditioning on *Z* the clusters). The underlying test performed by `ccdf` in the second DEA now consists in comparing the marginal CDF on *Z* with the conditional CDF on both *X* and *Z*. The two CDFs (the dotted line and the solid lines) are much closer which means the variable *X* has less impact on the conditional CDF. The number of steps of the CDF matches the number thresholds chosen in `ccdf` (10 in the analysis). The first value of each CDF is the proportion of zeroes.

An obvious limit of this real data analysis lies in the double use of the data. First Kusnadi *and others* (2021) performed a clustering to define cell populations that were not annotated beforehand. Then, we use once again those clusters of cells as a covariate in our DEA. The clustering is based on the cell gene expression, therefore the association between the clusters and the gene expression – needed for the clusters to play the role of a confounding variable – is constructed during the first step. Although this example was used as an illustration of the `ccdf` method, it is quite realistic thanks to high-throughput technologies that allow nowadays to measure cell characteristics (surface and intracellular proteins) as well as single-cell gene expression.

## 6. Discussion

We propose a new framework for performing CIT, with an immediate application to differential expression analysis of scRNA-seq data. Our approach can accommodate complex designs, e.g. with more than two experimental conditions or with continuous responses while adjusting for several additional covariates. `ccdf` is capable of distinguishing differences in distribution by using a CIT based on the estimation of CCDFs through a linear regression. The resulting asymptotic test is attractive due to the low computation times, especially dealing with a high number of observations. Yet, for small samples sizes (e.g. due to experimental design or cost limitations in data acquisition), we cannot always rely on an asymptotic test. Consequently, a permutation test is proposed in such cases. Performing permutations is obviously time consuming, but as it is necessary only for small sample sizes, computation times remain reasonable in such settings. Nevertheless, easy parallelization of the permutation test can alleviate this problem. Furthermore, an adaptive procedure, for both the number of permutations and the number of thresholds, is implemented in order to accelerate `ccdf` while preserving a sufficient statistical power along with numerical precision for the lowest *p*-values. Per-gene asymptotic tests can also be computed in parallel to speed up computations. The proposed approach has been fully implemented in the userfriendly R package `ccdf` available on CRAN at https://CRAN.R-project.org/package=ccdf.

While `ccdf` can be applied to many types of data thanks to its flexibility, it has been specifically tailored for the need of scRNA-seq data DEA. In the simulation study, `ccdf` exhibits great results and versatility in complex designs such as multiple conditions comparison, and also allows to analyze data when the experiment design includes confounding variables while most of the competing methods cannot. Finally `ccdf` maintains great power while ensuring an effective control of FDR. The application to a real single-cell RNA-seq data set enhances the importance of being able to handle complex experimental designs, especially with a potential confounding variable. Of note, `ccdf` being a distribution-free approach, it can support any normalization method (and this choice is ultimately left to the responsibility of the user).

Tiberi *and others* (2020) recently proposed non-parametric permutation approach that also compares empirical CDF. While `distinct` shares some common ideas with `ccdf`, the two approaches widely differs in their test statistics and capabilities. On the one hand `distinct` requires multiple samples and only addresses the two groups comparison, while on the other hand `ccdf` provides an asymptotic test and can accommodate more complex experimental designs.

The number of evaluating thresholds considered in `ccdf` directly impacts both its computation time but also potentially its statistical power. To optimize this trade-off and to maintain the performances while reducing the computational cost, we propose to use a sequence of evenly spaced thresholds. Besides, instead of choosing a regular sequence of thresholds, one can prefer to manually chose the different thresholds at which CCDFs are computed and compared, e.g. using *prior* knowledge to emphasize some areas of the distribution.

Although the linear regression estimation of the CCDF is efficient, different parameterizations can be considered (e.g. logistic regression which would be more natural). Currently `ccdf` cannot analyse multi-sample data, such as biological replicates or repeated measurements. This would consist in modifying the structure of the working model in equation (2.1). `ccdf` would thus be able to perform multi-sample analysis while keeping the advantage of its asymptotic test, at the cost of an increased computational burden. Besides, other test statistics could be investigated based on the same CIT.

## 7. Software

The proposed method has been implemented in an open-source R package called `ccdf`, available on CRAN at https://CRAN.R-project.org/package=ccdf and on Github at https://github.com/Mgauth/ccdf. All R scripts used for the simulations and the real data set analyses in this article were performed using R v3.6.3 and can be found on Zenodo at https://doi.org/10.5281/zenodo.5699462.

## Acknowledgments

MG is supported within the Digital Public Health Graduate’s school, funded by the PIA 3 (Investments for the Future - Project reference: 17-EURE-0019). The project is supported through SWAGR Inria Associate-Team from the Inria@SiliconValley program. Computer time for this study was provided by the computing facilities MCIA (*Mésocentre de Calcul Intensif Aquitain*) of the Université de Bordeaux and of the Université de Pau et des Pays de l’Adour. BH & MG thank Franck Picard for helpful discussion about this work.

