## Supplementary Material for "Distribution-free complex hypothesis testing for single-cell RNA-seq differential expression analysis"

**5 Application to a scRNA-seq study in COVID-19 patients** **12**

### 1 Method

#### 1.1 Parameters estimation

We have $\boldsymbol{Y} \in [\zeta_{\min}, \zeta_{\max}]$ for some known constants $\zeta_{min}, \zeta_{max}$. Let $\zeta_{\min} \leq \omega_1 < \omega_2 < ... < \omega_p < \zeta_{\max}$ is a sequence of $p$ ordered and regular thresholds. Let the design matrix $W$ be the matrix with $i$th row $\boldsymbol{W}_i = (1, \boldsymbol{X}_i, \boldsymbol{Z}_i)$ and $\tilde{\boldsymbol{Y}}_i = (\tilde{Y}_{11}, ..., \tilde{Y}_{1p})$ for the $p$ thresholds. The full set of regression coefficients for the $j$th regression is written $\widehat{\boldsymbol{\beta}}_j = (\widehat{\beta}_{0j}, \widehat{\boldsymbol{\beta}}_{1j}, \widehat{\boldsymbol{\beta}}_{2j})^\intercal$. By making use of OLS, we can write

$$\begin{aligned} \widehat{\boldsymbol{\beta}}_j &= (W^\intercal W)^{-1} W^\intercal \tilde{\boldsymbol{Y}}_j & (1)\\ &= (n^{-1} W^\intercal W)^{-1} n^{-1} W^\intercal \tilde{\boldsymbol{Y}}_j + o_p(1) & (2) \end{aligned}$$

where $\tilde{\boldsymbol{Y}}_j = (\tilde{Y}_{1j}, ..., \tilde{Y}_{nj})^\intercal$. Then,

$$\begin{aligned} \widehat{\boldsymbol{\beta}}_j &= n^{-1} \sum_{i=1}^{n} \Psi^{-1} \boldsymbol{w}_i \tilde{Y}_{ij} + \left\{ (n^{-1} W^\intercal W)^{-1} - \Psi^{-1} \right\} n^{-1} \sum_{i=1}^{n} \boldsymbol{w}_i \tilde{Y}_{ij} \\ &= n^{-1} \sum_{i=1}^{n} \Psi^{-1} \boldsymbol{w}_i \tilde{Y}_{ij} + o_p(n^{-1/2}) & (3) \end{aligned}$$

where $\Psi = \mathbb{E}(\boldsymbol{w}_i \boldsymbol{w}_i^\intercal) = \lim_{n\to\infty} n^{-1} W^\intercal W$. Denote $\boldsymbol{h}_i = (h_i^1, ..., h_i^s)^\intercal$ such as $\boldsymbol{h}_i$ is the matrix product of the rows of $\Psi^{-1}$ associated with $\boldsymbol{X}_i$ and $\boldsymbol{w}_i$. From (3), $\widehat{\boldsymbol{\beta}}_{1j}$ can be expressed by

$$\widehat{\boldsymbol{\beta}}_{1j} = n^{-1} \sum_{i=1}^{n} \boldsymbol{h}_i \tilde{Y}_{ij} \quad (4)$$

Recall that $\widehat{\boldsymbol{\beta}}_1 = (\widehat{\boldsymbol{\beta}}_{11}, \widehat{\boldsymbol{\beta}}_{12}, ..., \widehat{\boldsymbol{\beta}}_{1p})$ is a $s \times p$ matrix then we deduce from (4) the expression of the matrix of the $s$ coefficients corresponding to $\boldsymbol{X}$ for the thresholds $\omega_j$s, $\forall j = 1, ..., p$, as

$$\widehat{\boldsymbol{\beta}}_1 = n^{-1} \sum_{i=1}^{n} \boldsymbol{h}_i \tilde{\boldsymbol{Y}}_i \quad (5)$$

Last, in order to simplify, we transform the matrix $\widehat{\boldsymbol{\beta}}^1$ of size $s \times p$ into a vector $\widehat{\boldsymbol{\gamma}}^1$ of size $ps$ by concatenating the $s$ lines of $\widehat{\boldsymbol{\beta}}^1$ one after another.

Note that regression estimates are still consistent up to a multiplicative scalar [1], so the use of linear regression instead of logistic regression remains valid.

#### 1.2 Asymptotic test

After plugging in the estimate of $\beta_{1j}$ for the estimated test statistic, we need to derive the resulting asymptotic distribution under the null hypothesis. Knowing $\widehat{\boldsymbol{\beta}}_1$, by the multivariate central limit theorem, we have

$$\sqrt{n}\,(\widehat{\boldsymbol{\gamma}_1} - \boldsymbol{\gamma}_1{}^*) \longrightarrow N(0, \Sigma) \tag{6}$$

where $\boldsymbol{\gamma}_1^*$ is the true expectation of $\boldsymbol{h}_i\tilde{\boldsymbol{Y}}_i$ (concatenated into a vector). Under the null hypothesis, $\boldsymbol{\gamma}_1{}^* = \mathbf{0}$. $\Sigma$ is a symmetric positive semi-definite covariance matrix of size $ps \times ps$ that can be estimated by the method of moments: $\Sigma = n^{-1}\sum_{i=1}^{n}\mathrm{Cov}(\boldsymbol{h}_i\tilde{\boldsymbol{Y}}_i) = E\{\mathrm{Cov}(\boldsymbol{h}_i\tilde{\boldsymbol{Y}}_i)\}$ such as the $\big((k-1)p+j, (k'-1)p+j'\big)^{\mathrm{th}}$ entry of $\mathrm{Cov}(\boldsymbol{h_i}\tilde{\boldsymbol{Y}}_i))$ is defined as:

$$\begin{aligned}\mathrm{Cov}(\boldsymbol{h_i}\tilde{\boldsymbol{Y}}_i)_{(k-1)p+j,(k'-1)p+j'} &= \mathrm{Cov}\left(h_i^k \mathbb{1}_{\{Y_i \le \omega_j\}}, h_i^{k'}\mathbb{1}_{\{Y_i \le \omega_{j'}\}}\right)\\ &= \begin{cases} h_i^k h_i^{k'}\big(\pi_j - \pi_j\pi_{j'}\big), & \omega_j \le \omega_{j'} \\ h_i^k h_i^{k'}\big(\pi_{j'} - \pi_j\pi_{j'}\big), & \omega_{j'} < \omega_j \end{cases}\end{aligned}$$

where $\pi_j = P(Y_i \le \omega_j)$ for $j = 1, ..., p$ and $h_i^k$ is the $k^{th}$ element of $\boldsymbol{h}_i$ for $k = 1, ..., s$.

Let $\boldsymbol{\nu} = \sqrt{n}\widehat{\gamma}_1$, then

$$n\widehat{\gamma}_1^{\mathsf{T}}\widehat{\gamma}_1 = \boldsymbol{\nu}^{\mathsf{T}}\boldsymbol{\nu} = \boldsymbol{\nu}^{\mathsf{T}}\Sigma^{-\frac{1}{2}}\Sigma\Sigma^{-\frac{1}{2}}\boldsymbol{\nu} = \boldsymbol{u}^{\mathsf{T}}\Sigma\boldsymbol{u}^{\mathsf{T}} \tag{7}$$

where $\boldsymbol{u}$ follows asymptotically a multivariate standard normal. Let $\Sigma = UAU^{\mathsf{T}}$, where $U$ is an orthonormal set of eigenvectors and $A$ is a diagonal matrix of eigenvalues of $\Sigma$. Because of the orthonormality of $U$, $\tilde{\boldsymbol{u}} = \boldsymbol{u}U$ is also asymptotically multivariate normal.

By (6) and (7), the observed test statistic $\widehat{D}_n$ is given by:

$$\widehat{D}_n = n\sum_{j=1}^{ps}\widehat{\gamma}_{1j}^2 \tag{8}$$

and

$$n\sum_{j=1}^{ps}\widehat{\gamma}_{1j}^2 = \tilde{\boldsymbol{u}}^{\mathsf{T}}A\tilde{\boldsymbol{u}} = \sum_{j=1}^{ps}\widehat{a}_j\tilde{\boldsymbol{u}}_j^2 \tag{9}$$

which is asymptotically a mixture of $\chi_1^2$ random variables. The estimated mixing coefficient $\widehat{a}_j$ is the $j$th diagonal element of $A$. Note that the form of the estimated test statistic (8) is equivalent to the one given in the main manuscript.

Finally, by (9), we have the following asymptotic distribution of $\widehat{D}_n$ under the null hypothesis:

$$\widehat{D}_n \underset{n\to+\infty}{\longrightarrow} \sum_{j=1}^{ps}\widehat{a}_j\chi_1^2 \tag{10}$$

#### 1.3 Permutation test

Permutation tests are a simple way to obtain the sampling distribution for any test statistic, under the null hypothesis that there is no link between the outcome $Y$ and the variable $X$. The observations of $X$ can then be shuffled. Permutation tests are recommended when the number of observations is too small, so that the asymptotic distribution can not be assumed to hold. When the sample size $n$ is low, we propose to perform permutations to estimate the empirical distribution of $\widehat{D}_n$ under the null hypothesis. We distinguish two cases: i) testing the association between $Y$ and $X$ without any covariate and ii) testing the association between $Y$ and $X$ given a covariate $Z$.

**i) In the absence of covariates.** Under the null hypothesis, $Y$ and $X$ are independent, so the observations of $X$ are exchangeable. Hence, we can randomly permute the observations of $X$.

**ii) In the presence of covariates.** When we need to perform a conditional independence testing with a covariate $Z$, the observations of $X$ are not exchangeable without conditioning on $Z$. Indeed, if we randomly permute the observations of $X$, we break not only the link between $X$ and $Y$ but also the link between $X$ and $Z$. To preserve the dependency between $X$ and $Z$, we are facing two cases: (a) if $Z$ is a categorical variable and (b) if $Z$ is continuous. In case (a), we randomly switch $X$ within the groups defined by the categories of $Z$. Under scenario (b), the permutations become tricky. The idea is to permute two observations of $X$ only if the two corresponding observations of $Z$ are close. To do so, a conditional permutation algorithm based on the distance between the observations of $Z$ is proposed.

*Conditional permutation algorithm.* For now, the permutation test is only able to take into account an univariate variable $X$ and an univariate covariate $Z$ whereas the asymptotic test can handle a multivariate variable $X$ and a multivariate covariate $Z$ without increasing the computation times. Consequently, if one wants to adjust for many variables, the asymptotic test must be used. Even if the sample size is low, we show in the main manuscript that the asymptotic test remains powerful, so it is still reasonable to use this specific test in this case. When we need to adjust for a continuous covariate $Z$, the permutation test requires a specific shuffling. In order not to break the link between $X$ the variable to be tested and $Z$ the covariate, we permute the observations of $X$ according to a probability distribution $\mu_i$ which takes into account the relationship between $X$ and $Z$. $\mu_i$ is computed in the following way. First, we perform a linear regression of $X$ on $Z$, then for all $i = 1, ..., n$, for all $j \neq i$, we compute

$$\mu_{ij} = \frac{|\widehat{X}_i - \widehat{X}_j|^{-1}}{\sum_{\ell=1}^{n} |\widehat{X}_i - \widehat{X}_\ell|^{-1}} \tag{11}$$

with $\widehat{X}_i$ the predicted value coming from the linear regression. We denote

$\mu_i = (\mu_{i1}, \ldots, \mu_{in})$ the vector of probabilities computed according to (11). For a permutation step, the observation $X_i$ is then replaced by $X_i^\star$ randomly drawn from $\{X_\ell, \ell \neq i\}$ given the probabilities $\mu_i$. Therefore, for a given $i$, the observations close to $X_i$ given $Z_i$ have a higher probability to be chosen. By adding some randomness through $\mu_i$, $X_i$ is not replaced by the same observation at each permutation.

Following the appropriate method, we can permute the observations of $X$. Under i), we are able to compute the test statistic $D$ from the observations of $Y$ and the permuted observations of $X$ while under ii), the test statistic is obtained from the observations of $Y$ and $Z$ as well as the permuted observations of $X$. Then, under the null hypothesis, $B$ permutation-based test statistics $\{D_1^*, ..., D_B^*\}$ are calculated and $p$-values are computed as $\widehat{p} = \frac{1}{1+B}\left(1 + \sum_{b=1}^{B} \mathbb{1}_{\{\widehat{D}_n \leq D_b^*\}}\right)$, to avoid getting zero $p$-values [2], where $D_b^*$ is the test statistic obtained in the $b$-th permutation. Finally, we apply the Benjamini–Hochberg correction to the raw $p$-values to obtain FDR-adjusted $p$-values.

#### 1.4 Practical considerations for computational speed up

##### 1.4.1 Adaptive permutations

The disadvantage of using permutations could be the onerous computation times, especially when dealing with large sample size, which is more often encountered in single-cell DEA than in bulk DEA. The software computes 1,000 permutations by default for all the genes, but an adaptive procedure may provide similar accuracy at much lower computational cost. When calculation times appear to be too excessive, the user can switch to adaptive permutations. According to some pre-defined rules, the number of permutations is increased at each step to get sufficient numerical precision on the $p$-values only for certain genes. By default, the method computes 100 permutations for all genes, then we add 150 permutations for the genes with a $p$-value less than 0.1, bringing the total number of permutations for these genes to 250. Then, for the genes with an associated $p$-values less than 0.05, we perform 250 permutations more and finally the genes with a $p$-values less than 0.01, we add 500 permutations to reach 1,000 permutations for a reduced bunch of genes. If the computation times are still too long, the user can choose the number of $p$-values thresholds and the different limit values. The number of permutations executed at each step is also configurable.

##### 1.4.2 Evaluation thresholds

Selecting the thresholds $\omega_1, \omega_2, \ldots, \omega_p$ where the CCDF is evaluated may be difficult in practice. If too few thresholds are selected, then important changes in $F_{Y|X,Z}(\omega_j \mid x, z)$ may not be detected. One could instead select the thresholds to match the unique observations of $Y$, even though this selection technically violates the assumption that the thresholds are fixed and independent of the data. Yet, this technical violation does not appear to adversely affect the performance of our approach in simulations (see Section ??), and `ccdf` selects the thresholds to match the unique observations of $Y$ by default.

However, there may be an important computational cost to selecting so many thresholds: the number of linear regressions required to estimate all $\beta_1$s is then equal to the sample size. So when analyzing data from a large number of cells, one gets an equally large number of regressions to estimate along with a large matrix $\Sigma$, significantly increasing the computation time. One solution to reduce computation times (both for the asymptotic test and the permutation test) is to decrease the number of evaluated thresholds and thus the number of estimated $\beta_1$s as well as the dimension of $\Sigma$.

Instead of going through all the unique values of $Y$, one can choose a regular sequence of thresholds. Since single-cell RNA-seq data are count data, we propose spacing these thresholds according to a logarithmic scale, i.e., to better focus on the values where the CCDFs will not be too close to 1. This way, `ccdf` statistical power is maximized as variations in distributions are more likely to appear for smaller values (this point is all the more important as the number of thresholds is small).

### 2 Simulations

The code to generate the dataset under the described setting is available from the Zenodo archive [1]

#### 2.1 Analyses settings

`ccdf` We perform the asymptotic test with as many thresholds as unique values and without using a logarithmic threshold. The permutation test is computed using the adaptive procedure by default (from 100 to 1,000 permutations).

`SigEMD` We compute the Earth's mover distance to compare the distributions without using the imputation of dropouts (because the number of simulated dropouts is very low). `SigEMD`'s test performs a permutation test relying on two parameters to tune: the number of permutations and the bin size (between 0 and 1) to fix the width of the histogram's intervals. Given the fact the method exhibits huge computation times, we select the parameters with a good trade-off between the time of execution and the statistical power. Therefore, we arbitrarily fix the number of permutations to 500 and the bin size to 0.2.

`MAST` As described in the user guide [2], we fit a hurdle model and run a likelihood ratio test here, testing for differences in $X$.

[1] `https://doi.org/10.5281/zenodo.5699462`

[2] `https://www.bioconductor.org/packages/release/bioc/vignettes/MAST/inst/doc/MAITAnalysis.html#4_Differential_Expression_using_a_Hurdle_model`

`scDD` The method employs a Bayesian modeling framework requiring some hyperparameters. We refer to the method's quick start [3] and use the following settings: alpha=0.01, mu0=0, s0=0.01, a0=0.01, b0=0.01. We used the default option involving the Kolmogorov-Smirnov test. This allows to have a faster method despite a slight decrease of the power as indicated by the authors. We plan to add to the benchmark the full `scDD` framework based on permutations.

##### 2.2 2 conditions

Korthauer et al. [3] classified four different patterns of unimodal or multi-modal distributions:

- **differential expression (DE)**: two unimodal distributions with a different mean in each condition.
- **differential proportion (DP)**: two bimodal distributions with equal component means across conditions; the proportion in the low mode is 0.3 for condition 1 and 0.7 for condition 2.
- **differential modality (DM)**: one unimodal distribution in condition 1 and one bimodal distribution in condition 2 with one overlapping component. Half of the cells in condition 2 belongs to each mode.
- **both differential modality and different component means within each condition (DB)**: one unimodal distribution in condition 1 and 1 bimodal distribution in condition 2. The distributions have no overlapping components. The mean of condition 1 is half-way between the overall means in condition 2. Half of the cells in condition 2 belongs to each mode.

The non-differentially genes are divided into two categories:

- **(unimodal distribution with equivalent expression) (EE)**: two unimodal distributions with equal means.
- **bimodal distribution with equivalent proportions (EB)**: two bimodal distributions with equal component means across conditions and equal proportions in the low mode.

Since scRNA-seq data exhibits zero values (dropouts), we simulated beforehand a negative binomial distribution with mean equal to 0.5 in each condition to create zeroes or very low counts. Each distribution described above was simulated after uniformly having placed the zeroes and low counts between the 2 conditions, so that no difference in zero is created. With a sample size $n$,

[3] `https://bioconductor.org/packages/release/bioc/vignettes/scDD/inst/doc/scDD.pdf`

we chose a proportion of $n_0 = n/10$ for the zeroes and low counts equally divided between the conditions. The negative binomial distribution $NB(m,p)$ with size parameter equal to $m$ and probability parameter equal to $p$ has the following density: $\binom{x+m-1}{x}p^m(1-p)^x$ for $x \in \mathbb{N}^*$ and $p \in ]0,1]$. For simplicity, we kept constant $p = 0.5$ across all the scenarios. Then, for a sample size $n$, we denote $N = n - n_0$, gene expression is generated with the following parameters: $m_1 = \mathcal{U}(10,20)$, $m_2 = 3m_1$, $m_3 = m_1 + m_2/2$, $\alpha_0 = 0.5$, $\alpha_1 = n/3$ and $\alpha_2 = 1 - \alpha_1$.

- **DE**: for $i = 1,\ldots,250$,

$$y_{ij} = \begin{cases} NB(m_1,p) & \text{if } j = 1,\ldots,N/2 \\ NB(m_3,p) & \text{if } j = N/2+1,\ldots,N \end{cases}$$

- **DM**: for $i = 251,\ldots,500$,

$$y_{ij} = \begin{cases} NB(m_2,p) & \text{if } j = 1,\ldots,N/2 \\ \alpha_1 NB(m_1,p) + \alpha_2 NB(m_2,p) & \text{if } j = N/2+1,\ldots,N \end{cases}$$

- **DP**: for $i = 501,\ldots,750$,

$$y_{ij} = \begin{cases} \alpha_1 NB(m_1,p) + \alpha_2 NB(m_2,p) & \text{if } j = 1,\ldots,N/2 \\ \alpha_2 NB(m_1,p) + \alpha_1 NB(m_2,p) & \text{if } j = N/2+1,\ldots,N \end{cases}$$

- **DB**: for $i = 751,\ldots,1000$,

$$y_{ij} = \begin{cases} \alpha_0 NB(m_1,p) + (1-\alpha_0) NB(m_2,p) & \text{if } j = 1,\ldots,N/2 \\ NB(m3,p) & \text{if } j = N/2+1,\ldots,N \end{cases}$$

- **EE**: for $i = 1001,\ldots,5500$,

$$y_{ij} = NB(m_1,p) \quad \forall j = 1,\ldots,N$$

- **EB**: for $i = 5501,\ldots,10000$,

$$y_{ij} = \alpha_0 NB(m_1,p) + (1-\alpha_0) NB(m_2,p) \quad \forall j = 1,\ldots,N$$

#### 2.3 4 conditions

The four scenarios under 4 conditions comparison, inspired by Korthauer et al. [3], are the following:

- **multiple DE**: 4 unimodal distributions with a different mean in each condition.
- **multiple DP**: 4 bimodal distributions with equal component means across conditions; the proportion in the low mode is 0.1 for condition 1 and 0.3 for condition 2, 0.9 for condition 3, 0.7 for condition 4.

- **multiple DM**: 2 unimodal distributions in condition 1 and 2 and 2 bimodal distributions in condition 3 and 4 with one overlapping component. Half of the cells in condition 3 and 4 belongs to each mode
- **multiple DB**: 1 unimodal distribution in condition 1; 1 bimodal distribution in condition 2, 1 distribution with three modes in condition 3 and 1 distribution with 4 modes in condition 4; distributions in condition 2 and 4 have no overlapping components with distributions in condition 1 and 3; distribution in condition 3 have the middle component overlapping the distribution in condition 1. The mean of condition 1 is half-way between the overall means in condition 2, 3 and 4. Half of the cells in condition 2 belongs to each mode, a third of the cells in condition 3 belongs to each mode and a quarter of the cells in condition 4 belongs to each mode.

The non-differentially genes are divided into two categories:

- **multiple EE**: four unimodal distributions with equal means.
- **multiple EB**: four bimodal distributions with equal component means across conditions and equal proportions in the low mode.

Since scRNA-seq data exhibits zero values (dropouts), we simulated beforehand a negative binomial distribution with mean equal to 0.5 in each condition to create zeroes or very low counts. Each distribution described above was simulated after uniformly having placed the zeroes and low counts between the 4 conditions, so that no difference in zero is created. With a sample size $n$, we chose a proportion of $n_0 = n/10$ for the zeroes and low counts equally divided between the conditions. Then, for a sample size $n$, we denote $N = n - n_0$, gene expression is generated with the following parameters: $p = 0.5$, $m_1 = \mathcal{U}(10, 20)$, $m_2 = 2m_1$, $m_3 = m_1 + m_2/2$, $m_4 = 3m_1$, $\alpha_0 = 0.25$, $\alpha_1 = 0.1n/4$, $\alpha_2 = 1 - \alpha_1$, $\alpha_3 = 0.3n/4$, $\alpha_4 = 1 - \alpha_3$, $\delta = 0.9m_3$, $\gamma = 0.6m_3$ and $\epsilon = 0.3m_2$.

- **multiple DE**: for $i = 1, \ldots, 250$,

$$y_{ij} = \begin{cases} NB(m_1, p) & \text{if } j = 1, \ldots, N/4 \\ NB(m_2, p) & \text{if } j = N/4 + 1, \ldots, N/2 \\ NB(m_3, p) & \text{if } j = N/2 + 1, \ldots, 3N/4 \\ NB(m_4, p) & \text{if } j = 3N/4 + 1, \ldots, N \end{cases}$$

- **multiple DM**: for $i = 251, \ldots, 500$,

$$y_{ij} = \begin{cases} NB(m_2, p) & \text{if } j = 1, \ldots, N/4 \\ 1/2NB(m_1, p) + 1/2NB(m_2, p) & \text{if } j = N/4 + 1, \ldots, N/2 \\ 1/3NB(m_1, p) + 1/3NB(m_2, p) + 1/3NB(m_3, p) & \text{if } j = N/2 + 1, \ldots, 3N/4 \\ NB(m_1, p) & \text{if } j = 3N/4 + 1, \ldots, N \end{cases}$$

- **multiple DP**: for $i = 501, \ldots, 750$,

$$y_{ij} = \begin{cases} \alpha_1 NB(m_1,p) + \alpha_2 NB(m_2,p) & \text{if } j = 1,\dots,N/4 \\ \alpha_2 NB(m_1,p) + \alpha_1 NB(m_2,p) & \text{if } j = N/4+1,\dots,N/2 \\ \alpha_3 NB(m_1,p) + \alpha_4 NB(m_2,p) & \text{if } j = N/2+1,\dots,3N/4 \\ \alpha_4 NB(m_1,p) + \alpha_3 NB(m_2,p) & \text{if } j = 3N/4+1,\dots,N \end{cases}$$

- **multiple DB**: for $i = 751,\dots,1000$,

$$y_{ij} = \begin{cases} NB(m_3,p) & \text{if } j = 1,\dots,N/4 \\ 1/2NB(m_3+\delta,p) + 1/2NB(m_3+\delta,p) & \text{if } j = N/4+1,\dots,N/2 \\ 1/3NB(m_3-\gamma,p) + 1/3NB(m_3,p) + 1/3NB(m_3+\gamma,p) & \text{if } j = N/2+1,\dots,3N/4 \\ 1/4NB(m_3-2\epsilon,p) + 1/4NB(m_3-\epsilon,p) & \\ +1/4NB(m_3+\epsilon,p) + 1/4NB(m_3+2\epsilon,p) & \text{if } j = 3N/4+1,\dots,N \end{cases}$$

- **multiple EE**: for $i = 1001,\dots,5500$,

$$y_{ij} = NB(m_1,p) \quad \forall j = 1,\dots,N$$

- **mutiple EB**: for $i = 5501,\dots,10000$,

$$y_{ij} = \alpha_0 NB(m_1,p) + (1-\alpha_0)NB(m_2,p) \quad \forall j = 1,\dots,N$$

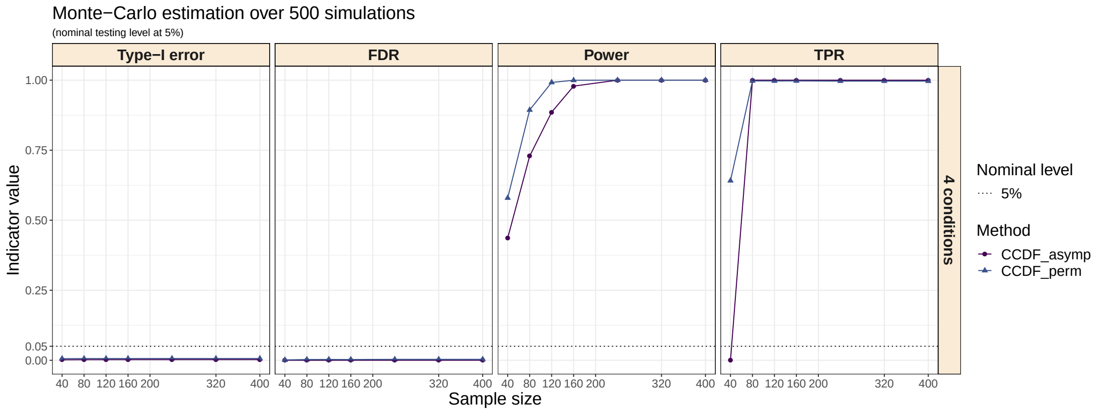

Figure 1: **Overall Type-I error, Power, FDR and TDR under the 4 conditions case with increasing sample size**.

#### 2.4 Two conditions comparison given a covariate $Z$

We simulated a confounding variable $Z$ from a Normal distribution $N(10,2)$. The variable to be tested $X$ was simulated in the following way:

$$X = \begin{cases} 1, & Z \le Q_1 \quad \text{and} \quad Q_2 \le Z \le Q_3 \\ 2, & \text{otherwise} \end{cases}$$

where $Q_p$ is the $p$th quartile of $Z$.

We generated Y as a normally distributed variable:

$$Y = \begin{cases} A * X + \epsilon_1, & \text{DE gene} \\ B * Z + \epsilon_2, & \text{non-DE gene} \end{cases}$$

where $\mu \sim N(1, 0.5)$, $A \sim N(5 + \mu \mathbb{1}_{\{X=1\}}, 1)$ , $B \sim U(0.3, 0.5)$, $\epsilon_1 \sim N(0, 1)$ and $\epsilon_2 \sim N(0, 0.5)$.

### 3 Comparisons using real data benchmarks

#### 3.1 Positive control dataset

Single-cell RNA-seq data from [4] are publicly available on GEO with the primary accession code GSE29087 for the positive control dataset. The top 1000 DE genes validated through qRT-PCR experiments and used as gold standard gene set [5] is available from the GitHub repository (https://github.com/Mgauth/ccdf). The matrix of raw counts contains 2,928 genes measured across 48 mouse embryonic stem cells and 44 mouse embryonic fibroblasts.

#### 3.2 Negative control dataset

Single-cell RNA-seq data from [6] are publicly available on GEO with the primary accession code GSE29087 for the negative control dataset. The matrix of raw counts includes 12,535 genes measured across 160 mouse embryonic stem cells.

#### 3.3 `ccdf` settings

As showed in the main document, the asymptotic test is powerful enough from a sample size of 80 observations. Furthermore, regarding the great number of genes, it is more appropriate to use the asymptotic test instead of the permutations due to the difference of computation times. The number of thresholds is equal to half the number of unique values for each gene.

### 4 Processing all types of data

`ccdf` is designed in the first place to perform single-cell DEA but the flexibility and the absence of distributional assumption on the input data allow to apply `ccdf` to any kind of variables as long as they remain continuous.

#### 4.1 Single-cell

The user must pay attention to two issues when dealing with single-cell measurements: the normalization and the dropouts.

##### 4.1.1 Normalization

`ccdf` does not contain a prior normalization step. The user must normalize the raw counts beforehand to make the samples comparable. Since there is no consensus on which normalization is most appropriate, the choice is left to the user and we advise to get acquainted with [7] for a comparative review.

##### 4.1.2 Dropouts

Single-cell data exhibit a huge amount of zeroes values. Even though `ccdf` is tailored to the presence or absence of zero inflation, if there is a large majority of zeroes (say more than 98%), one can question the accuracy of the results. We recommend to filter the genes by the number of dropouts that seems acceptable to the user and remove the selected ones from the analysis.

#### 4.2 Other types of data

Any kind of quantitative variables can be considered as an input of `ccdf` since no distributional assumption is made on the data. We propose the following example to show the versatility of `ccdf`. The user may not necessarily need to perform multiple tests, as in genomics data.

The Boston Housing Dataset was originally published by Harrison and Rubinfeld [8]. Each observation corresponds to one of the 506 neighborhoods near Boston along with 13 variables (see details here [4]). We want to test, say, the independence between per capita crime rate by town CRIM and median value of owner-occupied homes in $1000's MEDV, given lower status of the population LSTAT. Given the large sample size, we can use the asymptotic test (thanks to the corresponding function) with as many thresholds as unique values, since we perform a single test, and without logarithmic scale. If the sample size had been very low, a sub-function for the permutation test is also available.

### 5 Application to a scRNA-seq study in COVID-19 patients

[4] `http://www.cs.toronto.edu/~delve/data/boston/bostonDetail.html`

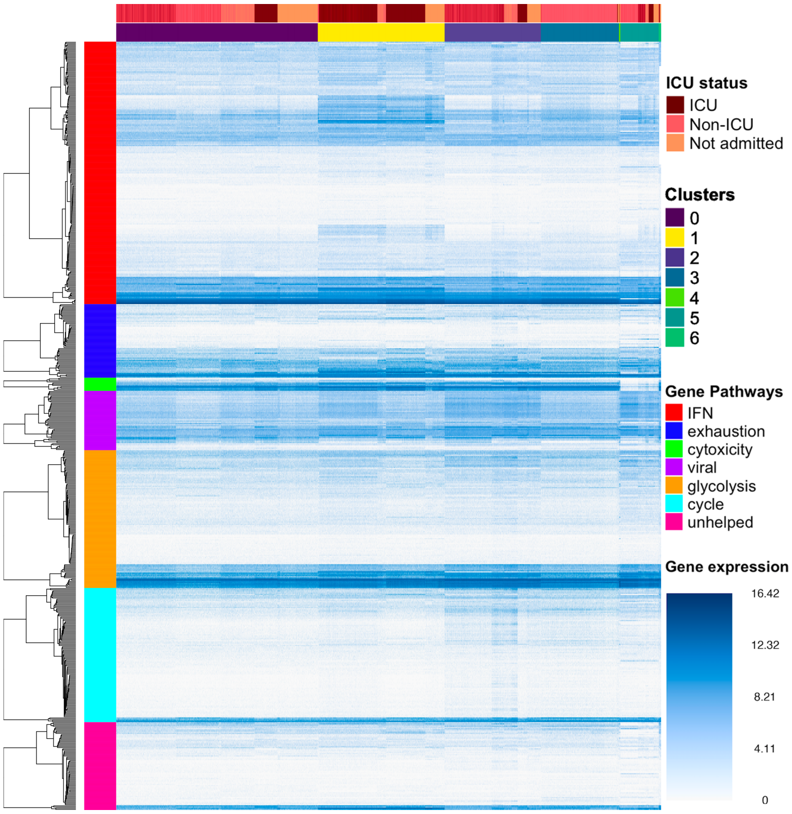

Figure 2: Heatmap of the log-CPM in the 7 gene pathways according to ICU status and clusters of cells.

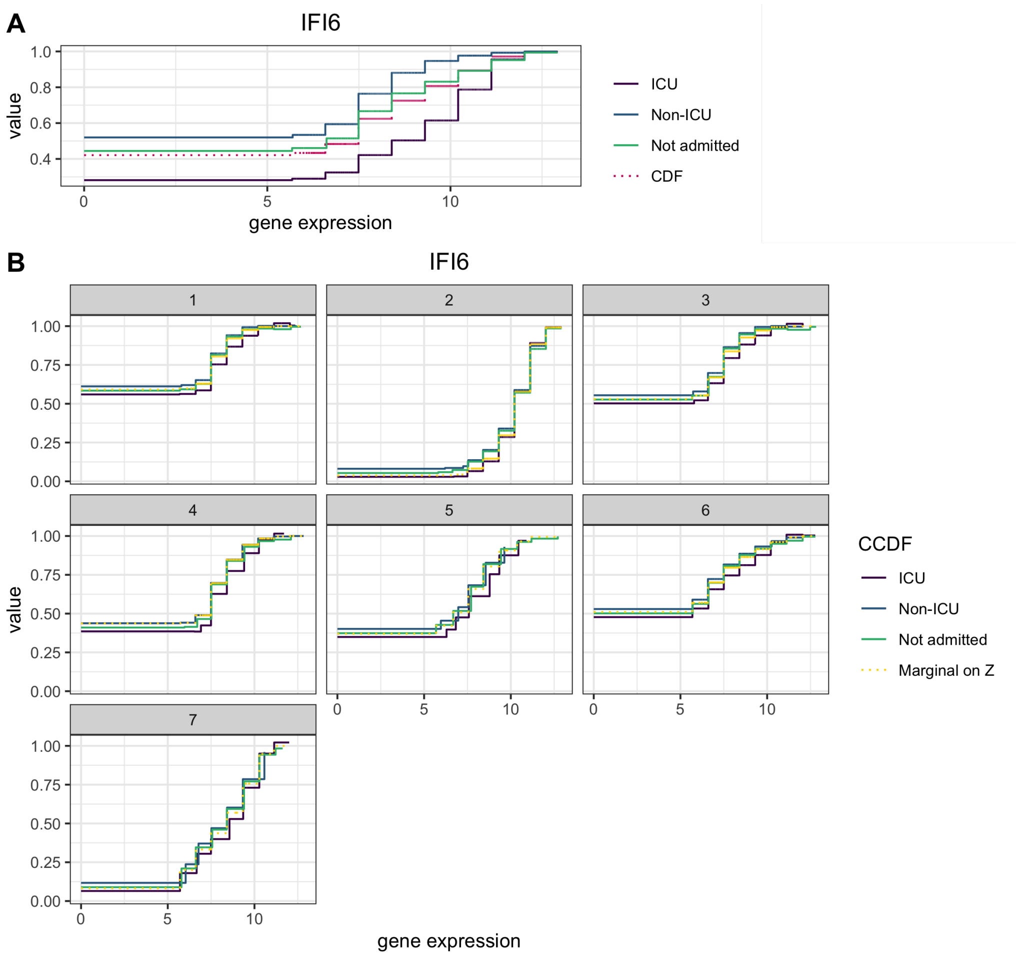

Figure 3: A: The solid lines represent the conditional CDF of IFI6 gene on ICU status (ICU, Non-ICU and Not admitted) and the dark pink dotted line represents the marginal CDF of IFI6 gene, *i.e.* without conditioning on ICU status. The underlying test performed by `ccdf` in the first DEA consists in comparing the marginal CDF with the conditional CDF. The $p$-value equals to 0 not adjusting for the clusters. B: The solid lines represent the conditional CDF of IFI6 gene on both $Z$, the 7 clusters, and $X$, the severity status (ICU, Non-ICU and Not Admitted), while the dotted yellow line represents the marginal CDF of IFI6 expression without conditioning on $X$ (but only conditioning on $Z$ the clusters). The underlying test performed by `ccdf` now consists in comparing the marginal CDF on $Z$ with the conditional CDF on both $X$ and $Z$. The two CDFs (the dotted line and the solid lines) are much closer which means the variable $X$ has less impact on the conditional CDF. The $p$-value equals to 6.204216e-36 when adjusting for the clusters. The number of steps of the CDF matches the number of thresholds chosen in `ccdf` (10 in the analysis). The first value of each CDF is the proportion of zeroes.

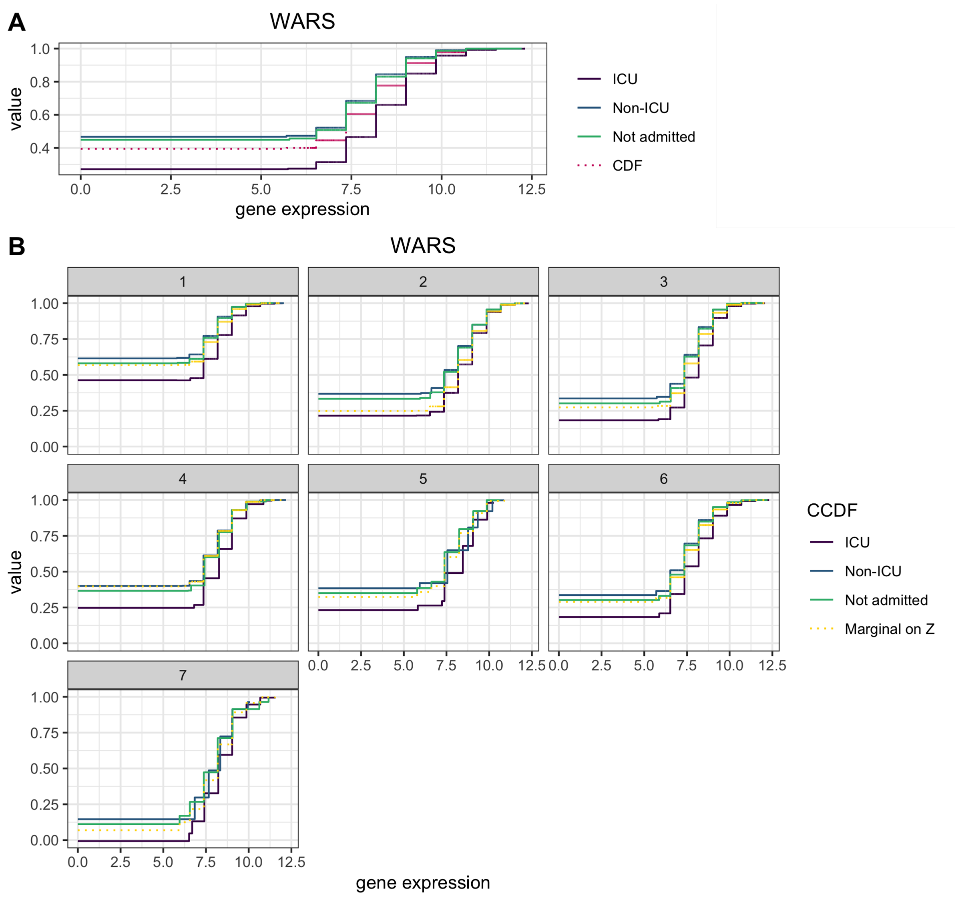

Figure 4: A: The solid lines represent the conditional CDF of WARS gene on ICU status (ICU, Non-ICU and Not admitted) and the dark pink dotted line represents the marginal CDF of WARS gene, *i.e.* without conditioning on ICU status. The underlying test performed by `ccdf` in the first DEA consists in comparing the marginal CDF with the conditional CDF. The $p$-value equals to 0 not adjusting for the clusters. B: The solid lines represent the conditional CDF of WARS gene on both $Z$, the 7 clusters, and $X$, the severity status (ICU, Non-ICU and Not Admitted), while the dotted yellow line represents the marginal CDF of WARS expression without conditioning on $X$ (but only conditioning on $Z$ the clusters). The underlying test performed by `ccdf` now consists in comparing the marginal CDF on $Z$ with the conditional CDF on both $X$ and $Z$. The $p$-value equals to 0.1347311e-201 when adjusting for the clusters. The number of steps of the CDF matches the number of thresholds chosen in `ccdf` (10 in the analysis). The first value of each CDF is the proportion of zeroes.

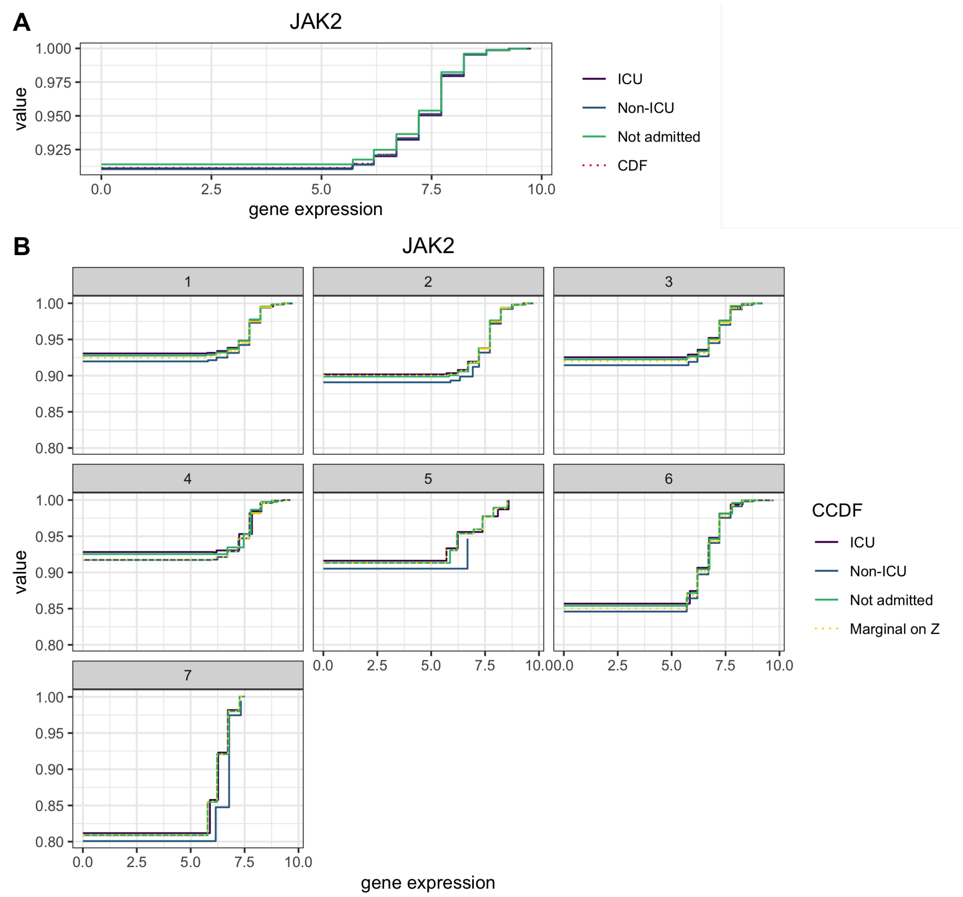

Figure 5: A: The solid lines represent the conditional CDF of JAK2 gene on ICU status (ICU, Non-ICU and Not admitted) and the dark pink dotted line represents the marginal CDF of JAK2 gene, *i.e.* without conditioning on ICU status. The underlying test performed by `ccdf` in the first DEA consists in comparing the marginal CDF with the conditional CDF. The $p$-value equals to 0.2793991 not adjusting for the clusters. B: The solid lines represent the conditional CDF of JAK2 gene on both $Z$, the 7 clusters, and $X$, the severity status (ICU, Non-ICU and Not Admitted), while the dotted yellow line represents the marginal CDF of JAK2 expression without conditioning on $X$ (but only conditioning on $Z$ the clusters). The underlying test performed by `ccdf` now consists in comparing the marginal CDF on $Z$ with the conditional CDF on both $X$ and $Z$. The $p$-value equals to 0.02076532 when adjusting for the clusters. The number of steps of the CDF matches the number of thresholds chosen in `ccdf` (10 in the analysis). The first value of each CDF is the proportion of zeroes.

[5] Moliner A, Ernfors P, Ibanez CF, Andäng M. Mouse embryonic stem cell-derived spheres with distinct neurogenic potentials. Stem cells and development. 2008;17(2):233–243.

[6] Grün D, Kester L, Van Oudenaarden A. Validation of noise models for single-cell transcriptomics. Nature methods. 2014;11(6):637–640.

[7] Lytal N, Ran D, An L. Normalization Methods on Single-Cell RNA-seq Data: An Empirical Survey. Frontiers in Genetics. 2020;11:41.

[8] Harrison Jr D, Rubinfeld DL. Hedonic housing prices and the demand for clean air. Journal of environmental economics and management. 1978;5(1):81–102.
